# Cytoskeletal tension actively sustains the migratory T cell synaptic contact

**DOI:** 10.1101/437236

**Authors:** Sudha Kumari, Michael Mak, Yehchuin Poh, Mira Tohme, Nicki Watson, Mariane Melo, Erin Janssen, Michael Dustin, Raif Geha, Darrell J. Irvine

## Abstract

When migratory T cells encounter antigen presenting cells (APCs), they arrest and form radially symmetric, stable intercellular junctions termed immunological synapses which facilitate exchange of crucial biochemical information and are critical for T cell immunity. While the cellular processes underlying synapse formation have been well-characterized, those that maintain the symmetry, and thereby the stability of the synapse remain unknown. Here we identify an antigen-triggered mechanism that actively promotes T cell synapse symmetry by generating cytoskeletal tension in the plane of the synapse through focal nucleation of actin via Wiskott -Aldrich syndrome Protein (WASP), and contraction of the resultant actin filaments by myosin II. Following T cell activation, WASP is degraded, leading to cytoskeletal rearrangement and tension decay, which result in synapse breaking. Thus, our study identifies and characterizes a mechanical program within otherwise highly motile T cells that sustains the symmetry and stability of the T cell-APC synaptic contact.

## Introduction

During an immune response, T cells form specialized junctions with cognate antigen presenting cells (APCs) termed ‘immunological synapses’ (Grakoui et al., 1999; Negulescu et al., 1996). Such synapses provide a stable platform where a number of T cell surface receptors including T cell receptors (TCRs), adhesion, and costimulatory receptors are enriched and engaged with counter-ligands on the APC surface (Dustin, 2008a; Miller et al., 2002; Negulescu et al., 1996). A symmetric, sustained synapse is a hallmark of T cell activation, both *in vitro* and *in vivo* (Bajenoff et al., 2006; Germain et al., 2012; Mempel et al., 2004; Miller et al., 2002; Stoll et al., 2002). It remains poorly defined how T cells – which are otherwise highly motile (Bousso and Robey, 2003; Miller et al., 2003; Tadokoro et al., 2006) – sustain their synaptic contacts with the APCs for prolonged durations. This is a crucial gap in our understanding of T cell biology since synapse lifetime is a critical determinant of T cell activation and function (Celli et al., 2007; Hugues et al., 2004; Skokos et al., 2007; Zaretsky et al., 2017).

Integrins, the actin cytoskeleton, and calcium signaling are all known to play important roles in the initial formation of an immunological synapse (Comrie and Burkhardt, 2016; Dustin et al., 1997; Fooksman et al., 2010; Martin-Cofreces and Sanchez-Madrid, 2018; Wei et al., 1999), but their roles in the subsequent maintenance and eventual dissolution of the synaptic contact are unclear. For example, integrin activation and actin polymerization are essential processes for T cell adhesion and synapse formation, but these processes could also terminate the synapse by engaging new adhesions and cellular protrusions, respectively, and thus driving T cell migration away from the APC. Furthermore, T cells are inherently highly migratory and have vigorous F-actin dependent lamellar undulations even in the synaptic phase (Roybal et al., 2013; Sims et al., 2007). These lamellar dynamics could promote polarization and lateral movement of the T cell (Mullins, 2010), thereby breaking the synapse symmetry. F-actin along with the activated integrin lymphocyte function-associated antigen-1 (LFA-1) is organized into a ring-like structure in the early synapse and this ring-like pattern is thought to promote synaptic junctional stability (Babich et al., 2012; Kaizuka et al., 2007; Wulfing et al., 1998). However, this F-actin/Integrin ring undergoes continuous fluctuations and therefore is amenable synapse symmetry breaking (Lomakin et al., 2015; Mullins, 2010; Sims et al., 2007). In addition, there exist genetic lesions in actin regulatory proteins such as the Wiskott-Aldrich Syndrome Protein (WASP), where T cells are able to adhere to and form synapses with APCs, but the resultant synapse are highly unstable (Calvez et al., 2011; Cannon and Burkhardt, 2004; Kumari et al., 2015; Sims et al., 2007; Thrasher and Burns, 2010). Thus, the role of the basic motility apparatus, especially the actin cytoskeleton, in sustaining T cell-APC contacts is complex and warrants further investigation.

Maintenance of the immune synapse is known to be influenced by T cell-intrinsic factors such as the strength of TCR signaling (Bohineust et al., 2018; Henrickson et al., 2008), however the downstream mechanisms at the level of the cellular migration machinery that enact the transition from an arrested to a motile state are not clear (Celli et al., 2007; Hugues et al., 2004; Shulman et al., 2014; Skokos et al., 2007). One of the ways in which the TCR-associated actin dynamics could enforce synapse symmetry and stability is by regulating mechanical forces within the synapse. Polymerization of actin is known to generate forces (Ridley et al., 2003), and organization of the polymerized F-actin network tunes the magnitude these forces (Blanchoin et al., 2014; Fletcher and Mullins, 2010). Notably, adhesion forces in T cell-APC conjugates are actin-cytoskeleton-dependent and continue to evolve after initial synapse formation (Bashour et al., 2014; Hosseini et al., 2009; Hu and Butte, 2016; Lim et al., 2011), raising the possibility that a specialized F-actin organization and associated forces may regulate maintenance of the synaptic contact.

We hypothesized that an examination of the actin cytoskeleton and associated forces in T cells during synaptic contact breaking would provide important clues to the mechanical design principles that T cells employ for maintaining the immune synapse. We examined T cell cytoskeletal organization at different stages of activation using a combination of super-resolution imaging, genetic and pharmacological perturbations, micromechanical measurements, and computational simulations, using both model APC-mimetic surfaces and physiological APCs. We found that specialized actin microstructures termed actin foci, which form within the interface following antigen encounter, generate and sustain intracellular tension within the T cell at the contact interface, and this tension actively sustains the synapse after its formation. This process relies on continuous nucleation of actin at TCR microclusters by Wiskott -Aldrich syndrome Protein (WASP), and interaction of freshly polymerized actin filaments with myosin II. Myosin contractile activity generates and maintains high in-plane tension across the synaptic interface. This high-tension actomyosin network eventually breaks as activated T cells downregulate WASP, leading to immediate relaxation of cytoskeletal tension followed by synaptic unraveling and resumption of motility. Taken together, these results uncover a novel antigen-triggered biomechanical program that regulates synapse symmetry in primary T cells, and highlight sub-synaptic actin architectures that sustain their interactions with APCs.

## Results

### Synapse maintenance during T cell activation is associated with actin foci but not integrin activation or calcium signaling

The actin cytoskeleton, integrins, and intracellular calcium flux all play important roles in mediating initial T cell motility arrest and formation of a symmetric synapse with antigen presenting cells. We thus first examined how these factors impact the maintenance of pre-formed synapses. To model synapse formation and eventual disengagement we seeded mouse primary naïve CD4^+^ primary T cells (referred to as ‘T cells’ henceforth) onto an anti-CD3/intercellular adhesion molecule-1 (ICAM-1)-coated coverslip (antigen presenting surface, APS) and allowed synapses to form. We then assessed cellular polarization over time (by recording the cells’ morphologic aspect ratio, AR) to detect T cells breaking their synaptic contact, because the contact interface shape is tightly linked to the motility state of the T cell (Hons et al., 2018; Houmadi et al., 2018; Mayya et al., 2018; Negulescu et al., 1996) (Figure 1A, Figure S1-2); Synapses are radially symmetric in their stable state, and polarize to break symmetry and acquire an elongated morphology prior to motility resumption (Videos S1-2).

**Figure 1.**
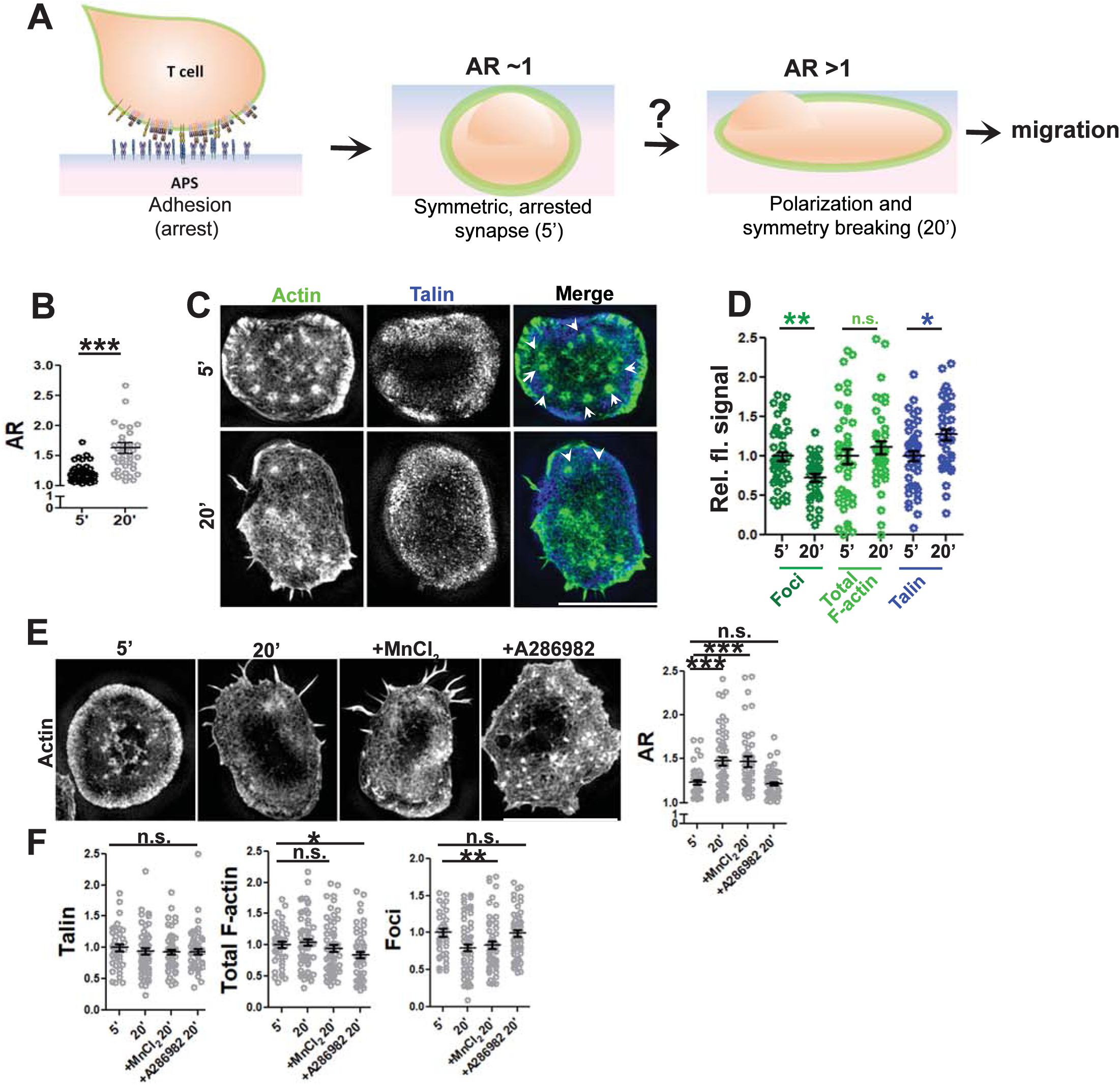
Synapse polarization involves actin remodeling in T cells and proceeds despite of integrin augmentation. (A) General schematic of the assay system used in this study for examining the process of synapse sustenance vs. polarization, post initial T cell adhesion and spreading on the antigen presenting surface (APS). Cell-APS interface of T cells was imaged for synapse establishment and maturation (5’) and eventual symmetry breaking and polarization (20’). (B-D) T cell interface actin re-organization during synapse symmetry breaking, imaged using SIM. (B) Quantification of the shape, or, relative fluorescence signal of the indicated proteins in the SIM images (c, d), normalized to levels detected at 5’; points represent values for individual cells. p values are: ** =0.003 for foci, **= 0.008 for talin, and p =0.36 for total F-actin using Mann-Whitney two-tailed test between populations of cells within the same experiment. (E-F) Integrin augmentation does not rescue synapse symmetry breaking. Cells were allowed to adhere to the APS for 5’ or 10’, and were then treated with vehicle control, 0.5mM MnCl2 or 100nM A286982 for subsequent 10’. Cells were fixed, stained and imaged using SIM (E) and analyzed for shape change (E) as well as talin recruitment, F-actin and foci at the synapse (F). p values in the graph, n.s. > 0.05; *** <0.001; *= 0.01; **<0.009 using Mann-Whitney two-tailed test between populations of cells within the same experiment. Scale bar, 5µm.

Within seconds, T cells seeded onto an APS spread to form a symmetric contact interface (Figure S1). Symmetric synapses persisted for ∼10 minutes, followed by a transition period when the cells began polarizing and becoming motile (Hons et al., 2018; Houmadi et al., 2018; Mayya et al., 2018; Negulescu et al., 1996), reflected in an increase in AR (Figure 1B, Videos S1-2). Structured illumination microscopy (SIM) revealed the presence of peripheral actin-rich lamellipodia and punctate actin structures distributed across the synaptic interface (Figure 1C) – termed ‘actin foci’ (Kumari et al., 2015)– that are formed at TCR microclusters upon TCR triggering. As T cells began to polarize at later times (20 min), actin foci were reduced while total F-actin levels in the interface remained constant (Figure 1C-D). Note that the foci quantification methodology we used (employing a Gaussian mask generated by a 1.6×1.6 µm^2^ rolling window, as described previously (Kumari et al., 2015) identified near-complete loss of foci as a ∼65% reduction in foci intensity, due to residual signal contributed by the lamellipodial F-actin (Figure S3). As T cells formed synapses with an APS, the key integrin adaptor protein talin accumulated rapidly at the periphery of the interface; talin levels remained constant at later times as cells began polarizing (Figure 1D). Activating integrins using MnCl_2_ (Dransfield et al., 1992) after a stable synapse was established did not affect subsequent polarization/synapse breaking at 20 min, and inhibiting LFA-1/ICAM interactions using the small molecule inhibitor A286982 (Liu et al., 2000) blocked T cell polarization, indicating that integrin activation is not necessary to sustain synapse symmetry and integrin activation may in fact be required for synapse symmetry breaking (Figure 1E). Notably, treatment of cells with A286982 was accompanied by a retention of actin foci at 20’, supporting a previous report that activated integrins may reduce F-actin accumulation in foci(Tabdanov et al., 2015). Under all of the treatment conditions, talin recruitment as well as the total F-actin was unchanged (Figure 1F left and middle graphs), however, higher actin foci levels were associated with reduced synapse polarization at late times (Figure 1F, right graph). Elevation of intracellular calcium is a key signal triggering initial arrest and formation of adhesive contacts between T cells and APCs (Miller et al., 2004). However, similar to integrin inhibition, blocking calcium signaling in T cells that had formed a synapse using BAPTA (Balagopalan et al., 2018) induced a retention of actin foci and synapse symmetry at 20’ (Figure S4B). Thus paradoxically, integrins and calcium signaling, which are critical for initial T cell arrest and formation of a symmetric synapse following TCR triggering, may also play roles in late polarization and synapse disruption. By contrast, actin foci were clearly enriched in the interface of T cells forming symmetric synapses and reduced or absent as T cells polarized and regained motility under all of these treatment conditions.

### T cell polarization is associated with WASP/actin foci downregulation

To directly visualize the dynamics of actin foci as T cells transitioned from the arrested to motile state, we imaged LifeAct GFP-expressing T cells using lattice light sheet microscopy (LLSM) – a technique that exposes cells to lower phototoxicity than traditional microscopy (Chen et al., 2014), allowing long term imaging of T cell synaptic actin dynamics that was not possible with confocal fluorescence imaging. LLSM imaging revealed that T cells engaged in a stable synapse continuously nucleate and dissipate actin foci across the contact interface, but as the cells polarized to begin migration, pre-existing foci were lost and new foci ceased to form (Figure 2A; Video S3). Thus, the presence of actin foci correlates with arrested synapses, and their loss accompanies synapse symmetry breaking.

**Figure 2.**
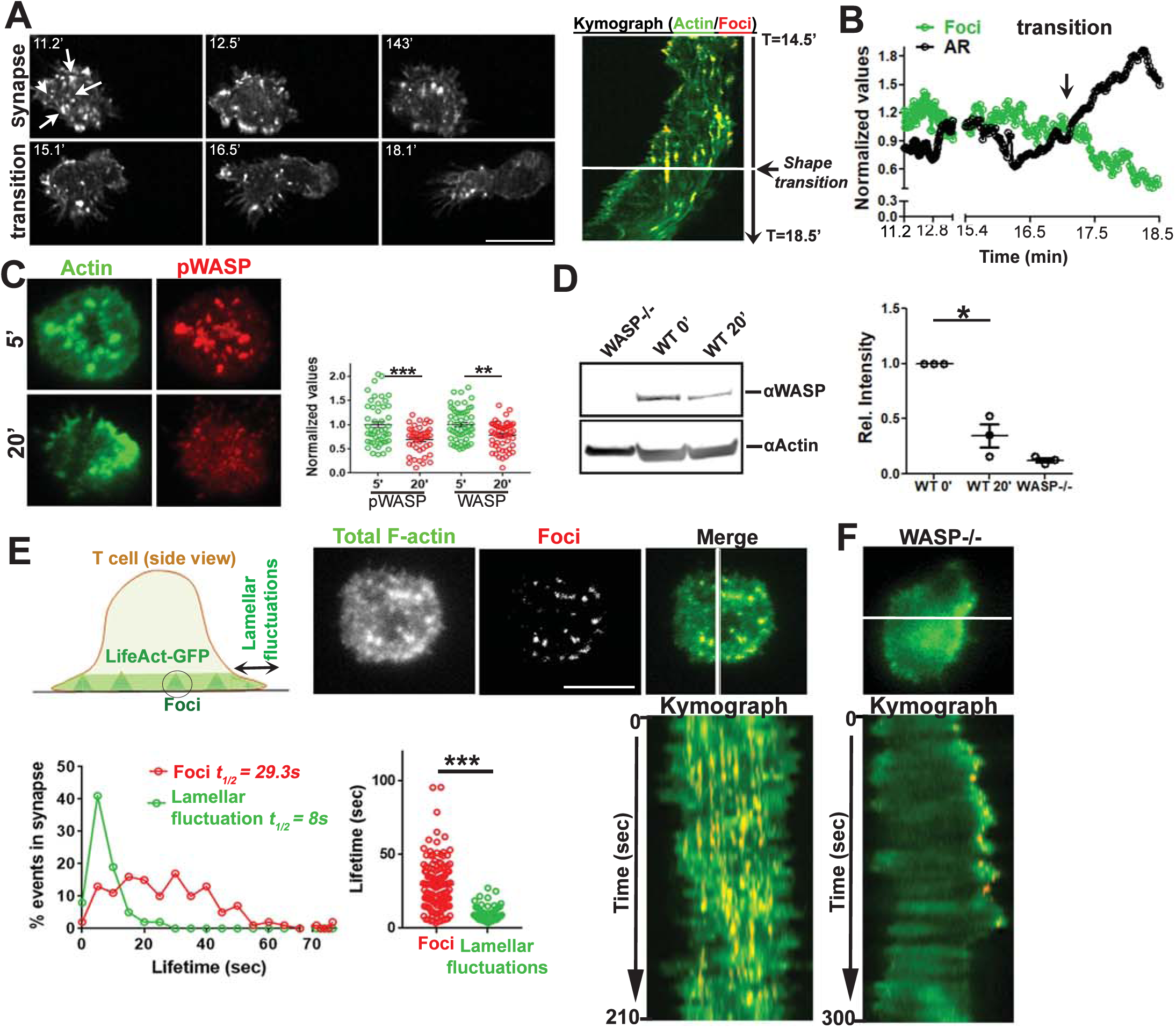
Synapse symmetry involves temporal regulation of WASP and WASP-dependent actin dynamics. (A) Time-lapse imaging of LifeAct-GFP expressing T cell using Lattice lightsheet microscopy (LLSM) during synapse polarization. Shown are snapshots at selected timepoints (left) and a kymograph of actin foci (denoted by arrows in the first image panel) over time (right) for a representative T cell. (B) Quantification of changes in actin foci and cellular aspect ratio in LLSM images during symmetry breaking. (C) T cells incubated with APS for the indicated time periods were processed for immunostaining (C) or for western blotting (D). Graph in (C) shows relative levels of total and active WASP (WASP-phosphorylated at Y293, pWASP), normalized to the mean levels at 5’. Numbers at the bottom of the western blot in (D) indicates WASP band intensity divided by the actin band intensity, and resulting ratios normalized to the values at the 0’ time point (non-activated cells). (E-F) Actin dynamics in mature synapse of LifeAct-GFP expressing T cell using TIRFM. The images show the foci (pseudocolored red) extracted from the total synaptic F-actin, and kymograph shows the time course of foci and lamellar activity over a period of 210 sec. Histograms of foci and lamellar protrusion and retraction (fluctuations) dynamics in WT T cell (E); Lifetime of individual foci and lamellar events (graph on the bottom right, each dot represents a single foci/lamellar protrusion-retraction event). ***<0.0001 using Mann-Whitney two-tailed test between populations of cells within the same experiment. (F) Shows a kymograph of the lamellar activity of WASP-deficient cells that lack actin foci. Scale bar, 5µm.

We posited that the loss of actin foci during synapse disengagement may result from WASP downregulation since the actin foci in the synapse are nucleated by WASP (Kumari et al., 2015) and TCR signaling-induced degradation of WASP has been reported previously (Macpherson et al., 2012; Reicher et al., 2012; Watanabe et al., 2013). Consistent with this idea, we detected a sizeable reduction in total as well as active forms of WASP (phospho-Y293) at 20’ post T cell activation in the T cell-substrate interface by TIRF microscopy as well as at the whole cell level by western blotting (Figure 2C,D). Simultaneously, highly dynamic lateral protrusions and retractions at the cell periphery (lamellar fluctuations) (Roybal et al., 2013) occur throughout the duration of a stable synapse (Figure 2E, Video S4). Comparison of foci and lamellar dynamics showed that foci had a mean half-life ï3 times longer than the lamellar actin fluctuations in WT cells (Figure 2E). In WASP^-/-^ cells lacking foci, lamellar fluctuations dominated the overall F-actin dynamics at the synapse (Figure 2F, VideoS4), as expected. Thus, even during the stable phase of the arrested synapse, actin polymerization drives constant lamellar fluctuations in the periphery that could drive symmetry breaking (Blanchoin et al., 2014; Carvalho et al., 2013), while actin foci are dynamically nucleated across the interface.

### Synaptic actin foci dynamics and associated traction forces support synapse symmetry

We previously demonstrated that actin foci are maintained via continuous nucleation of actin filaments at TCR microclusters (Kumari et al., 2015), a process that can generate localized mechanical stresses (Cossart, 2000; Pollard and Borisy, 2003). To directly quantify foci-dependent forces in the synaptic contact interface, we employed traction force microscopy (TFM) (Engler et al., 2004; Poh et al., 2012; Yeung et al., 2005): T cells were seeded on elastic polyacrylamide gels surface-functionalized covalently with anti-CD3 and ICAM-1 and containing embedded fluorescent beads. Forces exerted by the T cells on the substrate deform the substrate, leading to translocation of beads embedded in the gel from their resting positions (Figure 3A). Following 5 min of seeding onto the substrates, T cells were acutely detached by addition of EDTA and bead translation during substrate relaxation was measured to calculate forces exerted by the T cells. Using this approach, we found that WT T cells exerted substantial forces on the substrate (Figure 3A). These forces were greatly diminished in WASP^-/-^ cells lacking actin foci (Figure 3A).

**Figure 3.**
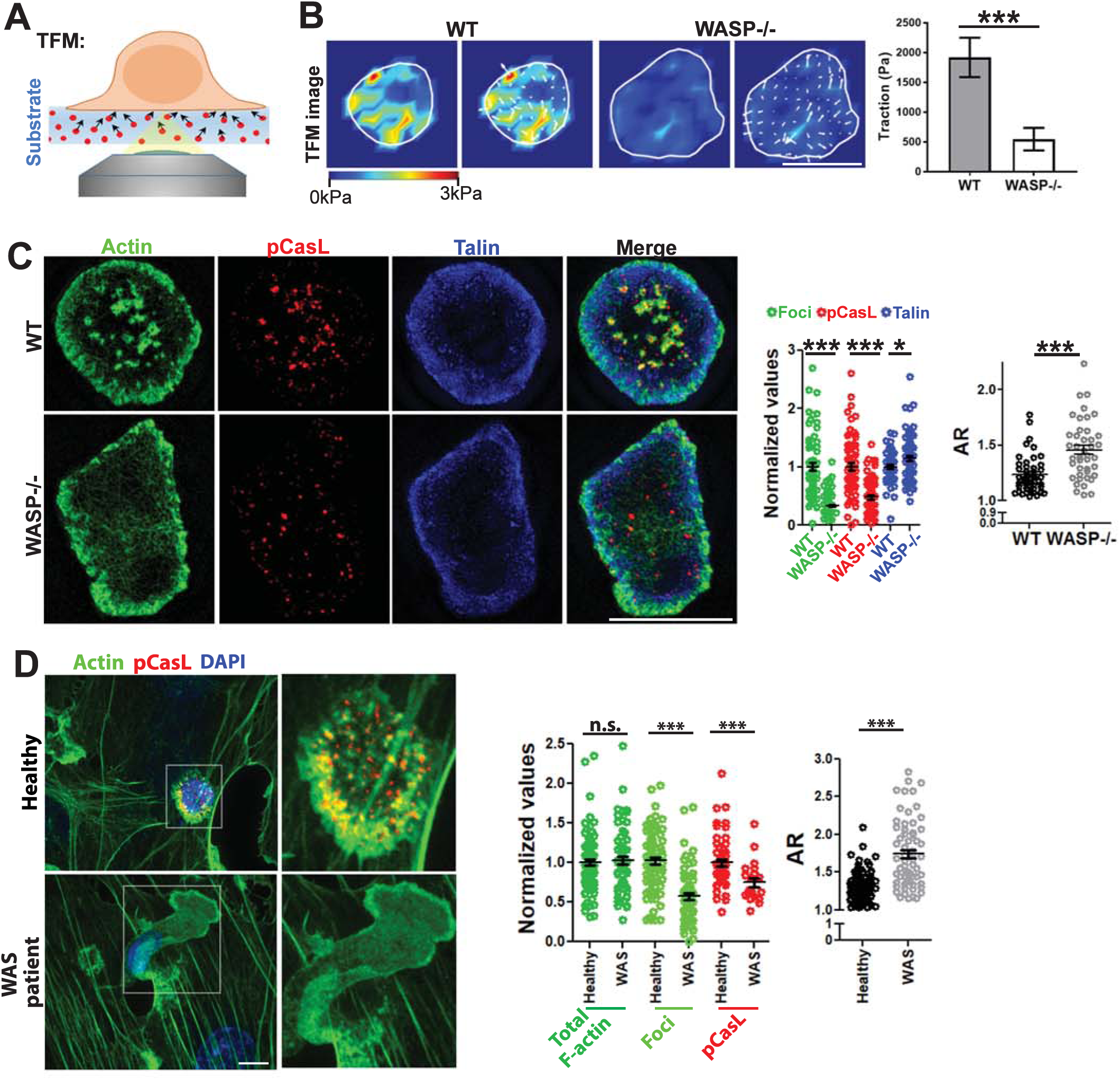
Foci associated mechanical forces are linked to synapse symmetry. (A-B) Foci –deficient cells display poor traction forces in their synapse. WT or WASP^-/-^ T cells were incubated on polyacrylamide substrates covalently functionalized with anti-CD3 and ICAM1, and traction force measurements were carried out as described in ‘Methods’. The images in the right show traction force maps without (left panels) or with (right panels) force vectors. p value, ***<0.0001. (C) SIM imaging of WT or WASP^-/-^ T cells activated by APS for 5’. The graphs show quantification of actin foci pCasL and talin levels in the synapse (Middle graph); right graph shows AR measurement *** *=*0.0001; p value for talin =0.07, as determined by using Mann-Whitney two-tailed test between populations of cells within the same experiment. (D) Human WAS patient CD4^+^T cells-APC conjugates display synapse symmetry defects similar to those observed in mouse WASP^-/-^ CD4^+^ T cells. CD4^+^T cells purified from healthy controls or WAS patients were incubated with superantigen loaded HUVEC cells for 5’ (APC; see ‘Methods’) and imaged using confocal microscopy. Each image represents a maximum intensity projection of the synaptic area. In the graphs, p values; *** <0.001; n.s. =0.09, as determined by using Mann-Whitney two-tailed test between populations of cells within the same experiment. Scale bar, 5µm.

The association of foci with mechanical stresses in the synapse was further reflected in their localization with the phosphorylated form of the mechanosensory protein CasL (pCasL). CasL is recruited to TCR microclusters in T cells upon antigenic stimulation, and undergoes a conformational change allowing phosphorylation of a cryptic N-terminal domain in response to local cytoskeletal stretching (Janssen et al., 2016; Santos et al., 2016; Sawada et al., 2006) (Figure S5A), which could be recognized by an antibody raised against pY249 p130Cas-a paralog of CasL. In activating T cells, pCasL significantly localized with a TCR signalosome marker-the phosphorylated form of Zap70 (pZap70) and with actin foci (Figure 3C, Figure S5B-C), and was reduced by the actin cytoskeleton-disrupting agent latrunculin A (Figure S5D), as expected for cytoskeletal tension-dependent phosphorylation of CasL. WASP^-/-^ T cells lacking actin foci showed lower levels of pCasL in the interface (Figure 3C), and at late times, polarizing WT T cells also showed lower pCasL levels (Figure S5E), indicating lower cytoskeletal stresses in these cells. Furthermore, even though WASP^-/-^ cells adhered and showed the same initial spreading kinetics and symmetrical morphology as WT T cells (Figure S6A), they formed unstable synapses with much earlier polarization and resumption of motility compared to WT cells (Figure S6B; VideoS5). The instability in WASP^-/-^ synapses was not due to grossly perturbed antigen receptor signaling: various key features of early TCR signaling and even the recruitment of CasL to the synapse were preserved in WASP^-/-^ cells (Figure S7). Further, unstable WASP^-/-^ synapses were also not due to reduced integrin signaling, as WASP^-/-^ cells did initially engage ICAM-1 in their synaptic interface and recruited normal amounts of talin (Figure S8). Furthermore, treatment with MnCl2 (Dransfield et al., 1992) (Figure S8) or supplementation of cytoplasmic Ca^2+^ levels using acute treatment with the thapsigargin (Gouy et al., 1990) (Figure S9) did not block the early polarization of WASP^-/-^ cells, indicating that integrin activation and calcium signaling are insufficient to reconstitute synapse symmetry in these cells.

To discount the possibility that lower mechanical stress in the interface and faster synapse symmetry breaking in WASP^-/-^ cells was an artifact of the minimal activation surfaces used here, or is a feature specific to murine T cells, we utilized CD4^+^ T cells derived from human WAS patient cells activated using live antigen presenting cells. Similar to the case of the murine cells (Figure S10A), WAS cells showed poor mechanotransduction, whether activated on anti-CD3/ICAM-1-coated coverslips (Figure S11B) or APCs (Figure 3D, Figure S11A), and broke synapses more frequently, even though the total F-actin content was normal. Faster synapse symmetry breaking in WAS cells was also not due to a developmental defect, since transient reduction in WASP levels in healthy human T cells using short-hairpin RNAs (shRNAs) (Kumari et al., 2015) yielded similar defects in pCasL and symmetry as seen with WAS cells (Figure S11A). Furthermore, the reduced synaptic pCasL in these cells was due to the loss of the actin nucleation-promoting factor (NPF) activity of WASP– overexpression of a WASP construct defective in NPF activity was indistinguishable from total WASP deficiency (Figure S11B). Together, these data indicate that Lack of foci nucleation underlies premature symmetry breaking in WASP^-/-^ T cells.

### In-plane cytoskeletal tension bolstered by actin foci dynamics promotes synapse stability

TFM measurements and pCasL accumulation showed that actin foci are associated with mechanical forces in the synapse. However, traction forces themselves do not explain how foci could promote synapse symmetry, since traction does not necessitate interface symmetry and has the potential to drive contact breaking and cell migration (Parsons et al., 2010). To further explore the relationship of cytoskeletal forces with synaptic interface symmetry, we next utilized a computational model of the actin cytoskeletal architecture and synaptic interface mechanics. We employed a reductionist Brownian dynamics model of actin dynamics at the cell-substrate interface (Kim, 2015; Mak et al., 2016). This model is based on the distinct behavior of actin foci and the peripheral lamella, the two distinct cytoskeletal networks observed at the synapse (Figure 2E), and incorporated three key differences observed between these actin structures: (i) we model foci as sites with higher actin nucleation rates than the lamella (as measured in (Kumari et al., 2015)), (ii) foci were treated as sites where actin could be immobilized (modeling association with TCR microclusters/integrins) with a lower local unbinding rate than at the lamella (due to the longer lifetime of foci vs. peripheral lamellar fluctuations), and (iii) foci were spatially placed across the synapse compared to the peripheral lamella (Figure 4A). The T cell contact interface was approximated as an 8 µm square, and a thin volume of the cytoplasm just inside the plasma membrane at the contact was modeled (500 nm thick), mimicking the submembrane actin-rich region (Fritzsche et al., 2013; Linsmeier et al., 2016) (Figure 4A). Actin foci were modeled as a regular array of 400 nm square sites across the contact plane (small orange squares in Figure 4A), and the peripheral lamella was modeled as a 400 nm wide zone at each edge of the contact (green); Actin nucleation and actin-linked receptor adhesions to substrate ligands proceed in all of these zones (Figure 4A). In this framework, F-actin evolves through a series of reactions comprising basic elements of actin dynamics: polymerization/depolymerization/nucleation of actin filaments, actin crosslinking proteins linking/unlinking actin fibers, binding and unbinding of actin filaments to adhesion sites on the substrate, and myosin-mediated contractile forces applied to the cytoskeletal network, with actin nucleation and unbinding rates that differ between each zone.

**Figure 4.**
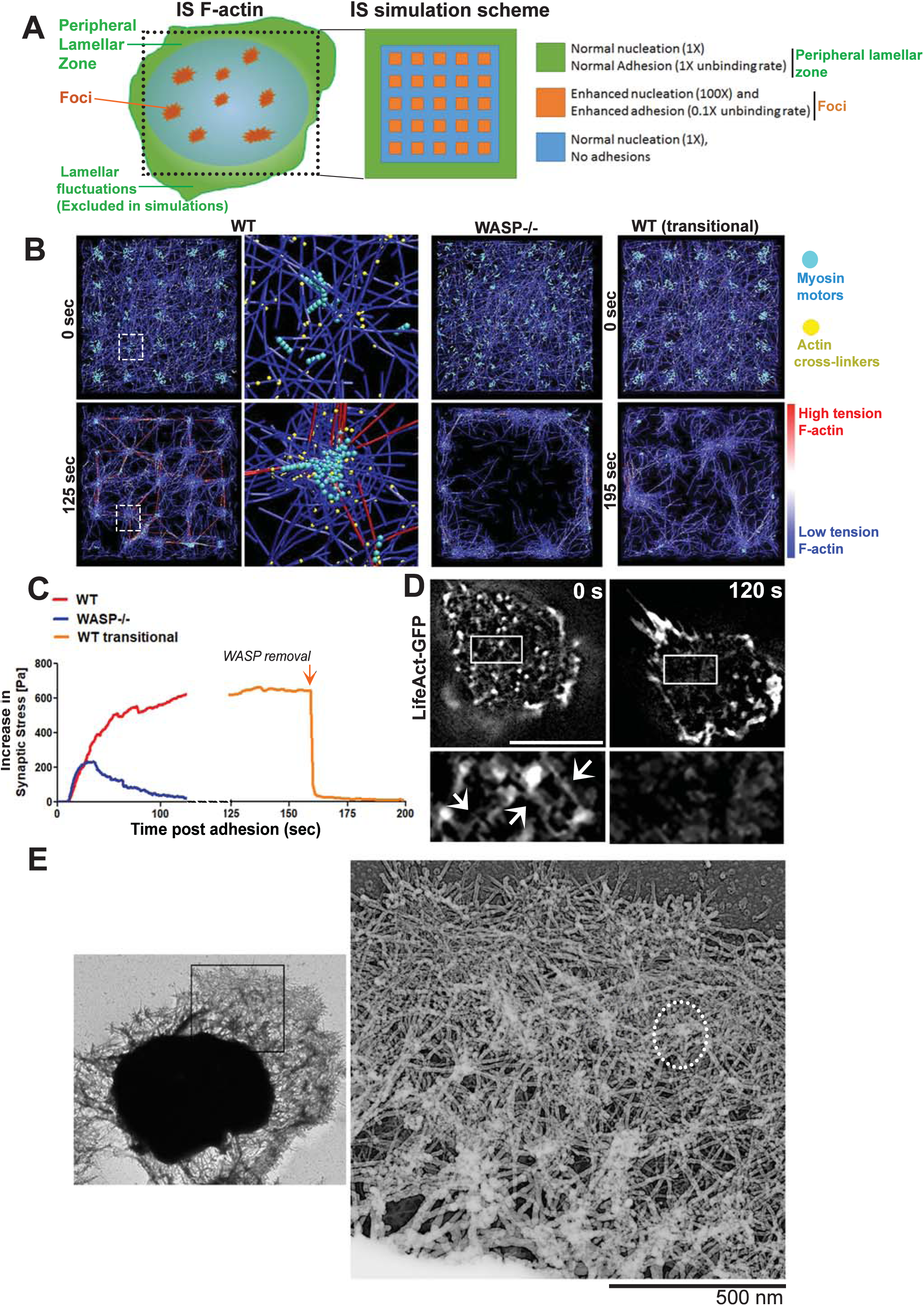
Computational simulations to explain the cytoskeletal basis of symmetry sustenance at synapse. (A) Outline of the spatial scheme utilized to simulate synapse cytoskeletal dynamics. Cartoon on the left shows the spatial patterning of the two major actin networks: the lamellar F-actin network in the peripheral zone that would continuously undergo fluctuations, and foci. The foci are interspersed throughout the synapse within a dynamic lamellar F-actin network. The peripheral lamellar fluctuations seen in live imaging could not be included in the simulations due to unavailability of data on fresh boundary adhesions. The cartoon on the right shows square simulation space patterned after the cartoon on the left and utilized here for simulations. The actin nucleation and adhesion parameters used at the foci and lamellar zones are indicated on the right. (B) Simulation of T cell IS F-actin behavior and resultant mechanical tension incorporating the differential dynamics and positioning of foci and lamella across the synaptic interface, using Brownian dynamics equation with no inertia (see ‘Methods’). The images show simulation snapshots at the beginning (0 min; soon after the attachment and spreading of T cells on the substrates) and the end of the simulations for the three IS cases – with persistent foci (WT, left most panels), with no foci (WASP^-/-^, middle right panels), and with WASP and consequently foci downregulated after a period of synapse maturation (WT transitional). The colors in the images indicate myosin II motor (pseudocolored teal), F-actin crosslinking proteins (pseudocolored yellow), and the F-actin filaments (pseudocolored blue for the low tension and the red for thigh tension actin filaments); middle left image panels show the distribution of the aforementioned proteins at individual foci site at a higher magnification. (C) Predicted stress (tension normalized to the area) profiles within the cytoskeletal architecture as the synaptic contact progresses in time, corresponding to the three cases shown in (D) Live imaging of a T cell using SIM shows the inter-foci connections (arrows in the inset) as predicted in the simulations; and their loss concomitant with loss of actin foci, as the T cell polarizes to initiate motility (compare insets between 0 and 120 sec). ‘0 sec’ refers to the beginning of the observation of the cell, after it has attached to the substrate, spread and maintained synapse for 12min. Scale bar, 5µm. (E) Ultrastructural visualization of T cell synaptic cytoskeleton using electron microscopy. The micrograph shows actin foci (identified as globular cluster of short filaments in the inset)-dependent interconnected cytoskeletal architecture that is associated with synapse symmetry as predicted in the model.

Simulations of synaptic actin dynamics, cytoskeletal network morphology, and the resultant network tension profile over time were examined under conditions when actin foci are present (modeling wildtype cells; ‘WT’) or absent (modeling WASP^-/-^ cells). In WT cells, these simulations revealed that actin foci acted as nodes supporting an interconnected actin network across the synaptic interface (Figure 4B left panels, Videos S6-7). This foci-connected actin network generated and sustained tension across the interface for prolonged durations (Figure 4C; synaptic stress is a cross-sectional area-normalized measure of the cytoskeletal network tension, parallel to the cell-substrate interface; see ‘Methods’), fueled by continual nucleation of filaments at the foci regions and associated myosin contractility (Videos S6-7). In the absence of actin foci (WASP^-/-^), peripheral lamellae dominate and the F-actin network largely accumulates in a lamella-like distribution (Figure 4B; Video S8), accompanied by a relaxation in overall synaptic cytoskeletal tension (Figure 4B,C). Modeling the late stages of the T cell synapse by first allowing a foci-rich network to form and then downregulating WASP (foci site associated actin nucleation), we saw that a rapid dissolution of foci and rapid redistribution of stable foci-anchored actin network to peripheral lamellar zones, coincident with attenuation of cytoskeletal tension (‘WT transitional’ Figure 4B-C, Video S9).

SIM imaging of F-actin dynamics in the near-interface region of the cytoplasm of live T cells undergoing the transition from the arrested to polarized state confirmed the macroscale changes in the F-actin network suggested by these simulations. In the stable synapse of a WT T cell expressing LifeAct-GFP, inter-foci connections were visible (Figure 4D ‘0 sec’). Both foci and their actin interconnections were lost as the cells polarized and regained motility (Insets, Figure 4D). These results suggest that when foci are present, a restraining tension within the actomyosin network is sustained and supported by foci, across the contact area. Without foci, F-actin is depleted from the central zone and largely enriched in the peripheral lamella, along with myosin activity, as expected in adherent cells poised to break symmetry (Lomakin et al., 2015; Yam et al., 2007). It is important to note that while it is the persistent lamellar protrusion that eventually would likely drive motility when symmetry is broken and synaptic stress is low (Lomakin et al., 2015; Yam et al., 2007), we could not model the protrusions in our simulations since experimentally measuring boundary adhesion dynamics in primary T cells is technically limited. Together, these simulations suggest that actin foci serve as nodes for tension generation within the synapse, which may contribute to their role in sustaining synapse symmetry.

### Synaptic tension required for synapse stability is generated by an interplay of actin foci and myosin II activity

In the computational model, myosin II contractile activity played a major role in generating cytoskeletal network tension across the synapse. Since local intracellular cytoskeletal tension dynamics are difficult to measure experimentally, we instead tested the role of myosin-mediated tension generation via acute pharmacological perturbations of myosin II to uncouple actin foci from myosin-mediated tension generation. Inhibition of myosin II using Blebbistatin (Blebb.) led to reduced pCasL levels and premature T cell polarization, even though foci and talin accumulation in the interface were not reduced (Figure 5A-C). Thus, myosin II functions downstream of foci to generate cytoskeletal contractile tension, and loss of this mechanical tension promotes T cell polarization. pCasL levels in Blebb.-treated cells were not as low as in WASP^-/-^ cells, which may reflect residual cytoskeletal stresses, potentially protrusive, arising from actin polymerization at the actin foci (Kumari et al., 2015; Pollard and Borisy, 2003). Blebb. treatment also did not further exacerbate the tendency of WASP^-/-^ T cells to break symmetry, as expected in our model (Figure 5D-E). Cells treated with the Arp2/3 complex inhibitor CK666 that preferentially ablates foci (Kumari et al., 2015), were included as a control, demonstrating that pharmacological inhibition of actin foci results in reduced pCasL and an increased tendency for T cell polarization (Figure 5B-D, Video S10). The modest effect of CK666 on contact symmetry could be due to the perturbation of lamellipodial actin protrusion required for symmetry breaking.

**Figure 5.**
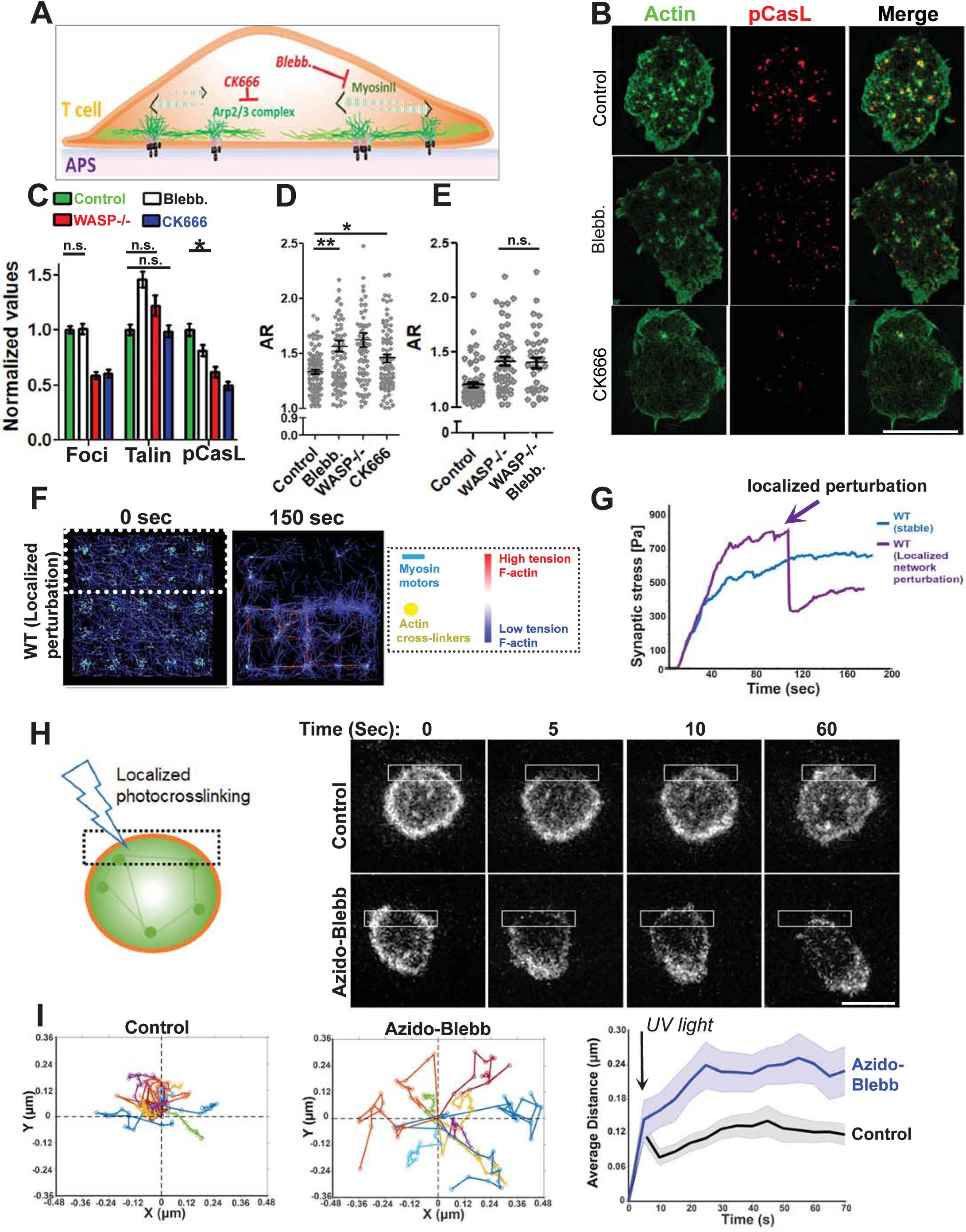
Pharmacological interrogation of actin foci-myosin crosstalk in sustaining symmetry. (A) Schematic of the cytoskeletal inhibitors and their targets used in the experiments. (B-E) SIM imaging of mouse T cells activated for 5’ on APS, then treated with the indicated inhibitors or vehicle control. Shown are the representative SIM images (B), and quantification of actin foci, talin, or pCasL in the contact interface (normalized to control cells, C), and measured cell morphology aspect ratios (D, E). Since CK666 treatment reduces F-actin intensity (Kumari et al., 2015), the ‘actin’ image of the CK666 treated cell in (B) was contrasted differently than the ‘control’ and ‘Blebb.’ cases for a clearer visualization of F-actin. p values, * =0.02, ** =0.001, n.s. >0.05. For the rest of the comparisons between the control and treatment cases, p <0.0001. For the bar graph in (B), at least 83 cells were used in each case; *p value* for the n.s. cases is 0.05, * =*0.02*. Within the foci, pCasL and talin groups, wherever not shown, the values between the Control and treatment is <0.0001 using Mann-Whitney two-tailed test between populations of cells within the same experiment. Scale bar, 5µm. (F-G) Simulation of synaptic F-actin network following localized myosin inactivation and inactivated myosin-mediated F-actin cross-linking within the area marked with dotted line in the left image. The graph shows the predicted alteration in mechanical stress profile, as myosin is inactivated crosslinked in the defined region (marked by an arrow), (G). (H-I) Experimental confirmation of localized actomyosin crosslinking and symmetry breaking. LifeAct-GFP expressing T cells were incubated with activating substrates, treated with vehicle control or 1µM Azido-blebb and their contact interface imaged using low excitation laser power (0.5 mW), using a spinning disc confocal microscope (H). The montage shows the shape change and movement that synapse undergoes after it is exposed to a brief flash of blue light crosslinking Azido-Blebb and actomyosin network within the illuminated area (shown with a box in the image). The deflection in center of mass (COM) within a minute was calculated and plotted for at least 10 cells in each case (graphs in H). The left graphs in (I) show individual COM trajectory, and the graph on the right shows average COM movement across cells within a minute post photo-inactivation of myosinII and network crosslinking. Scale bar, 5µm.

To further examine if the synapse-wide transmission of foci-dependent cytoskeletal tension is important for maintenance of a symmetric synapse, we carried out simulations of the actin network as in Figure 3, where we allowed the actin network to evolve for 120 seconds, and then selectively ablated F-actin network connectivity and tension in a spatially controlled manner (dashed area in the simulation snapshot of Figure 5E). This was achieved using localized myosin II inactivation and crosslinking of associated F-actin, and this perturbation was found to cause a disruption in synaptic F-actin organization and symmetry, as well as a significant lowering of overall synaptic tension (Fig. 5F-G, Video S11). To experimentally test the prediction that the symmetry of F-actin tension is important for synapse maintenance, we utilized a photoreactive form of Blebbistatin (azido-blebb) to optically crosslink the actomyosin network in a spatially controlled manner (Kepiro et al., 2012). Azidoblebb. binds to myosin II, and following excitation with UV light crosslinks the actomyosin network, effectively freezing actomyosin contractions in a spatially-controlled fashion. LifeAct-GFP-expressing T cells forming a symmetric synapse were treated with low doses of azido-blebb, photoactivated locally on one edge of the cell (dashed boxes in Figure 5H), and imaged under low excitation power to avoid phototoxicity. Photoactivation of the inhibitor induced an immediate deflection of the cellular center of mass and transition to a polarized morphology (Figure 5H-I, Video S12). Thus, symmetry maintenance requires tension transmission across the interface, mediated by high filament nucleation at actin foci and myosin contractile activity. The perturbation of this tensional mechanism via loss of foci, myosin II inhibition, or localized actomyosin crosslinking leads to a loss of cellular symmetry (Figure 6).

**Figure 6.**
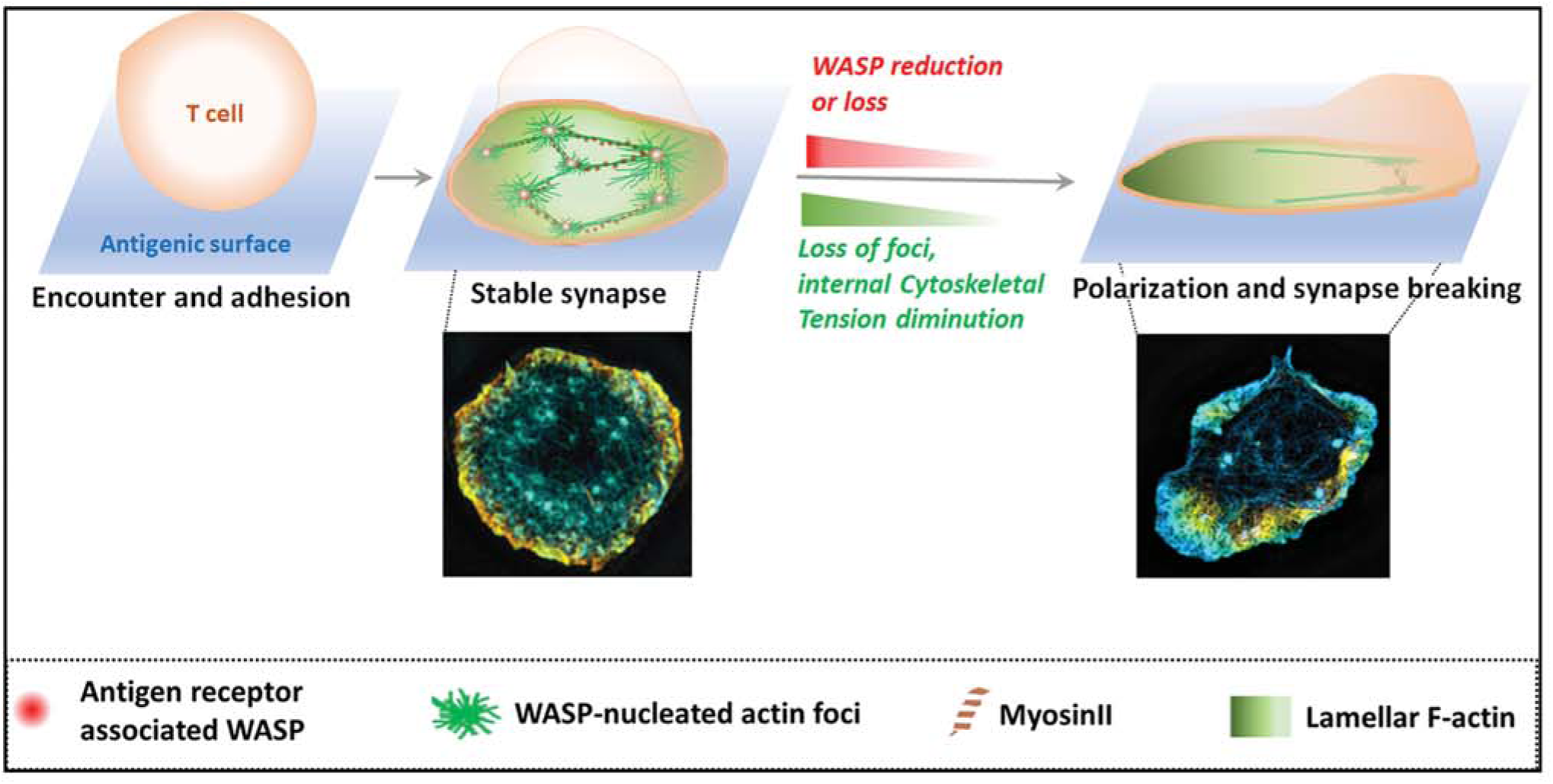
Model outlining the mechanistic basis of T cell synapse maintenance. The model argues that in highly migratory T cells, after adhesion and synapse formation, lamellar fluctuations would constantly attempt to break the synapse. An antigen-induced actin foci and associated actomyosin arrangement would generate high in-plane cytoskeletal tension that actively restrains synapse breaking. Via nucleation of actin foci, WASP acts as a central regulator of this mechanism. Downregulation of WASP following T cell activation leads to loss of foci-dependent actin architecture and internal tension resulting into synapse unravelling, as is also seen in WASP –deficient cells that are predisposed to synapse symmetry breaking. Images at the bottom of the cartoon show representative SIM images of F-actin at synapse.

## Discussion

Transitions in motility states underlie T cell immune function. The fast motility state of T cells is associated with immunosurveillance, while antigen-induced arrest is considered to be a hallmark of activation, both in *in vitro* and *in vivo* (Celli et al., 2007; Hugues et al., 2004; Mempel et al., 2004; Skokos et al., 2007; Zaretsky et al., 2017). Here, using naïve primary T cells, we have identified and characterized an antigen-induced cytoskeletal tension program that actively prolongs their arrested or ‘synaptic’ state. Loss of this mechanism, which normally happens when cells are sufficiently activated, leads to unraveling of the arrested synaptic contact followed by resumption of cellular motility. The experiments presented here reveal the cytoskeletal architectures that lymphocytes utilize to tune mechanical tension within the synapse in order to achieve a rapid shift in motility. These results also further our understanding of the mechanical deficits underlying deficient T cell function in Wiskott-Aldrich syndrome and provide a conceptual framework to assess synaptic organizational defects in other actin-related primary immunodeficiency diseases.

The process underlying immediate arrest of T cells when they encounter APCs displaying high-affinity antigens and concomitant formation of a symmetric contact has classically been termed the “stop signal” (Downey et al., 2008; Dustin, 2002, 2008b; Dustin et al., 1997; Hugues et al., 2004; Mempel et al., 2004). We find that actin foci are an integral part of the cellular machinery associated with maintaining TCR signaling-induced motility arrest. Previous studies have associated intracellular calcium flux in T cells with the stop signal (Dustin et al., 1997; Negulescu et al., 1996). While it is intuitive that actin foci machinery would communicate with the calcium stop signal pathway for coordinating synapse stability, it is unclear how this communication might take place. At the cellular level, we find evidence that the calcium pathway depends on WASP-dependent actin foci to promote synapse stability. A rise in intracellular calcium induced by thapsigargin is able to freeze shape transitions only in migrating WT T cells (data not shown). Furthermore, foci-lacking WASP^-/-^ T cells that show impaired intracellular calcium flux (Kumari et al., 2015) do not arrest when treated with thapsigargin. In contrast, we show that the foci-mediated synapse retention is independent of intracellular calcium because BAPTA treatment did not reduce foci or promote synapse symmetry breaking. Detachment of T cells from APCs was shown to be associated with a period of TCR signaling unresponsiveness (Bohineust et al., 2018); whether a reduction in TCR –induced calcium signaling during the transition from stable to motile synaptic interface might play a role in antigen unresponsiveness remains to be investigated. It is important to note that the methodology we used to examine the foci-intracellular calcium connection using pharmacological calcium elevation can generate minor off-target effects on contact symmetry and actin foci, since they alter some basic features of the synapse (e.g. talin downregulation in thapsigargin treatment). Furthermore, a previous study proposed that T cells could selectively utilize actin-based versus calcium-based mechanisms of synapse stability based on antigen affinity (Moreau et al., 2015). This implies that the calcium or foci-mediated stop-signal pathways could be selectively dominant depending on the strength of the TCR-antigen interaction.

Even though the foci, being F-actin-nucleating structures, might generate protrusive forces perpendicular to the interface, we find that the protrusive behavior of the foci orthogonal to the substrate may not be required for lateral synapse stability. Foci-mediated synapse stabilization proceeds regardless of the substrate stiffness (data not shown) and operates even when the substrate is too hard for the foci to protrude into (e.g. glass coverslips). Furthermore, inhibition of myosin predisposes synapses to break under conditions where foci nucleation and thus protrusion is not inhibited. Instead, our data suggests that actin foci act as stakes anchoring a tension-bearing actin network parallel to the interface plane (Figure 6). Myosin activity generates symmetric cytoskeletal tension throughout this network which we hypothesize counters the protrusive forces of the peripheral lamellae. The anchoring function of foci dominates over lamellar protrusions in part via their longer lifetime, and by virtue of foci being distributed across the synaptic interface. When these two requirements are met, actin foci can feed the actomyosin tensional network and stabilize synapses, irrespective of the substrate stiffness. When T cells are activated by supported lipid bilayers, actin foci are not static but move with TCR microclusters (Kumari et al., 2015). Yet in this system, the stability of synapse still relies on the presence of foci (Sims et al., 2007). The dependence of synapse stability on cell-intrinsic cytoskeletal dynamics rather than the substrate stiffness raises an exciting possibility that T cells could utilize cytoskeletal tension to sustain their synapse, when encountering antigens displayed by APCs of varying stiffness.

Myosin II plays a central role in foci-mediated synapse symmetry maintenance. The acto-myosin architectures have been characterized in lymphocytes previously (Babich et al., 2012; Carisey et al., 2018; Murugesan et al., 2016). A substantial fraction of myosin II is associated with formin-dependent actin arcs within the synapse of Jurkat T cells (Murugesan et al., 2016). The distinct architecture and positioning of the inter-foci actomyosin bundles described in primary naïve T cells in this study suggest that they are unlikely to be arcs (Figure 4). Whether the arcs themselves are involved in stabilizing the synapse and how they may communicate with the foci-actomyosin system to co-ordinate synapse stability remains to be investigated. Similarly, another study recently reported actin “vortice –like foci” that are interconnected by the actomyosin network in NK cells (Carisey et al., 2018; Fritzsche et al., 2017). It remains unclear whether the NK cell “vortices-like foci” cytoskeleton reported previously and the foci described here have related functions or involve similar molecular mechanisms.

At the functional and architectural level, actin foci represent a structure intermediate between podosomes and focal adhesions (FAs) described for innate immune cells and adherent cells, respectively. The continual actin nucleation at foci as well as some of their molecular and organizational characteristics resemble those of podosomes observed in innate immune cells (Cervero et al., 2018). Both are WASP-dependent, enriched in cortactin homolog protein HS1, are maintained by continuous actin nucleation, are associated with contact symmetry breaking in macrophages (Cervero et al., 2018) and are capable of generating protrusions. However, several key features distinguish the two: actin foci are antigen receptor-triggered (Kumari et al., 2015), and display a lifetime at least an order of magnitude shorter than the podosomes. Podosomes in macrophages show polarized localization during contact interface symmetry breaking (Cervero et al., 2018), while the foci in primary T cells show bulk downregulation during synapse symmetry breaking. Additionally, unlike podosomes, we predict that the protrusive behavior of foci is not required for synapse stability. In fact, the generation of filaments by foci to assemble an actomyosin system that organizes and scales mechanical tension across the contact interface reported here resembles the focal adhesions (FA) reported previously in fibroblasts (Kanchanawong et al., 2010). However, there is a key difference between the two mechanisms as well. FAs attach to stable actin filaments. Consequently, a cell can detach from the substrate by generating high forces that overcome tension at the FA and rupture them. In contrast, there is continual nucleation and generation of actin filaments at foci. Under these circumstances, forces of detachment would dissipate at foci and the only way to overcome the foci is to downregulate their formation, which is what we see in cells that are preparing to migrate. In this regard, the foci are a departure from the classical “clutch mechanism” proposed for the FAs (Kanchanawong et al., 2010), and represent an adhesion modality that T cells have evolved for their specialized motility requirements.

The results presented in this study indicate that mechanisms of synapse formation and synapse maintenance are mechanistically distinguishable. While increased F-actin is required for the former, the latter constitutes only a change in F-actin organization and overall F-actin content is constant. Similarly, activation of the integrin LFA1 is a prerequisite for successful synaptic engagement, but is not needed for maintenance of the synapse. These results support a recent finding that the LFA1-ICAM1 interaction is not necessary for stable CD4 T cell-DC synapses in the lymph node (Feigelson et al., 2018). The physiological implications are that T cells may use integrin-based adhesion to initiate synaptic contacts with selective cellular partners (Feigelson et al., 2018; Zaretsky et al., 2017), but maintain adhesion using lateral tensional mechanisms within the cell provided by the dynamic cytoskeletal arrangements comprising foci nucleation and acto-myosin contractions described here.

In their lifetime, T cells encounter antigens in diverse mechanical microenvironments. The antigen-induced tensional pathway identified here could represent a T cell-intrinsic tool to accommodate mechanical disparities between antigen presenting cells, and achieve optimal antigen sampling time. Different T cell subtypes are known to display varying motilities even when they encounter the same APC type (Tadokoro et al., 2006), and T cells encounter antigens of varying affinities for their TCRs. How the actin foci-based mechanisms outlined here operate in these diverse contexts remains an open question. Simultaneous examination of the effects of antigen affinity on foci, TCR triggering, and T cell activation in different T cell classes will help establish a relationship between TCR antigen affinity and mechanotransduction via this tensional mechanism at the immunological synapse. Future studies will also be needed to clarify the ways in which the described mechanical deficits in the synapses formed by WASP deficient T cells contribute to the WAS immunodeficiency at the whole-organism level (Thrasher and Burns, 2010).

## Acknowledgements

We thank A. Thrasher and D. Cox for WASP constructs, and J. Burkhardt and Nathan Roy for LifeAct-GFP mice. We thank the Advanced Imaging Center at Janelia Campus and J. Haddleston for help with the Lattice Light Sheet microscopy, and the Keck microscopy facility at the Whitehead Institute. We are thankful to F. B. Gertler and A. K. Dhawale for input on the experiments and data analysis. This work was supported in part by the Koch Institute Support (core) Grant P30-CA14051 from the National Cancer Institute. We thank the Koch Institute Swanson Biotechnology Center for technical support, specifically Eliza Vasile for OMX Deltavision microscope. This work was supported by the Ragon Institute of MGH, MIT, and Harvard, the National Institutes of Health grant AI43542 (MLD) and Wellcome Trust grant PRF 100262 (MLD), support from Yale University to MM and NIH grant 5R01AI100315 to RG. DJI is an investigator of the Howard Hughes Medical Institute.

## Methods

### Cells and reagents

For most experiments, unless otherwise mentioned, mouse primary CD4^+^ T cells were isolated from C57BL6/j WT or WASP-/-Mice lymph organs, using EasySep™ Mouse CD4+ T Cell Isolation Kit (StemCell). Cells were rested at 37°C for few hours in culture media (RPMI+ 10%FBS+ 100µM β-merceptoethanol + 1mM Sodium Pyruvate), and used for experiments thereafter. For live imaging, cells were cultured in phenol red free XVivo-10 media (Lonza) supplemented with 10units/ml IL-2. For antigen-specific CD4^+^T cells, CD4^+^ T cells were isolated from OTII WT or WASP-/-animals, using the procedure as described above.

For isolating human CD4+T cells, human peripheral blood from healthy donors was acquired from the New York Blood Bank, or from WAS patients at Boston Children’s hospital (two patients, males). The blood samples were and transported, handled and processed in accordance with the OSHA guidelines. The cells were isolated from the blood using the CD4^+^ rosette sep. kit, rested at 37°C in culture media for a few hours and were used in the experiments thereafter.

Unless otherwise mentioned, the inhibitors were purchased from Sigma chemicals, and were used at a concentration indicated in the figure legends. The antibodies for immunostaining including phopho-Y249Cas (#4014), phospho-Y171 Lat (#3581), phospho-Y397HS1 (clone D12C1), phosphor-Y139/SykY352 Zap70 (clone 65E4) were purchased from Cell Signaling Technology. Anti-phospho-Y145 (#EP2853Y, Novus), anti-tubulin (F2168, Sigma), anti-MyosinII (PRB440P, Covance), anti-WASP (Chicken polyclonal for immunostaining, #Gw22608, Sigma), anti-WASP (Rabbit polyclonal for immunostaining, #SAB4503087, Sigma), anti-phospho Y290 WASP (#Ab59278, Abcam), and anti-talin (clone C20, Santacruz), were purchased from the indicated vendors.

### Cell activation substrates

In most experiments glass coverslip (#1.5 chambered coverglass, Thermo Scientific) coated with 10ug/ml anti-CD3 and 1 µg/ml ICAM1 were used as activating substrate. For cell-cell conjugate experiments, either IFNγ (100units/ml, overnight)-treated and superantigen loaded (TSST-1+ SEB cocktail, 1µg/ml for 1hr at 37°C) HUVEC cells (Kumari et al., 2015), or antigen-loaded, differentiated primary BMDCs were used for synapse assays with human or mouse T cells respectively. In experiments examining the distribution of ICAM1 on activating substrates, planar lipid bilayer system was used. Glass-supported lipid bilayers were deposited and were reconstituted with anti-CD3-Alexa568, and ICAM1X12-his-cy5, and incubated with T cells as described previously (Kumari et al., 2015).

### Microscopy

#### Interference Reflection Microscopy

For IRM, LabtekII chambers were coated with anti-CD3 and ICAM1, were washed and incubated with the CD4^+^T cells suspended in the growth media. Care was taken to maintain the temperature of the T cells to 37°C, between the transfers from the culture dish to imaging chambers. The chambers were imaged using a temperature controlled 37°C stage adaptor on a Zeiss LSM510 confocal microscope equipped with a 63X 1.40 NA objective. The samples were imaged using the 543nm Laser at 0.1mW, and collected using Zeiss LSM software. Images were analyzed using Fiji Software.

#### Structured illumination microscopy

For SI superresolution microscopy, fixed cells were imaged with an OMX-3D microscope, V3 type (Applied Precision, GE), equipped with 405-, 488-, and 594-nm lasers and three Photometrics Cascade II EMCCD cameras. Images were acquired with a 100×, NA 1.4 oil objective at 0.125-µm Z steps using 1.518 immersion oil at room temperature. The images were acquired under the same illumination settings across the individual experiment and then processed with OMX SoftWoRx software (Applied Precision, GE). Images were saved in the TIFF format after reconstruction and alignment using optimized OTFs and wiener filter settings. The images reconstructed using SIM SoftWoRx processing software and were then analyzed using Image J.

#### TIRF

For the TIRF experiments, a Nikon revolution spinning disc confocal system equipped with a TIRF module and Andor iXON EMCCD camera was used. The images were acquired using 405nm, 488nm, 561nm, and 642nm lasers, a 100X objective (NA 1.49) and a 1.5X projection lens, and analyzed using Fiji software. The depth of TIRF field was maintained to 120nm-200nm, as measured by optically resolved fluorescent beads.

#### Confocal Microscopy and azidoblebbistatin experiment

For imaging cell-cell conjugates, Nikon revolution spinning disc confocal microscope equipped with Yokagawa CSU-X1 module was used. Images were acquired using 100X objective (NA 1.49) along with a 1.5X projection and an Andor iXON EMCCD camera. For azidoblebbistatin treatment experiment, the above mentioned spinning disc system was used to acquire a 0.4um optical section in the plane of the synapse, coupled with live microscopy. Cells were allowed to form synapses, and were treated with either inhibitor or DMSO just before imaging. During imaging, the inhibitor was activated by excitation with 5 brief pulses of 405nm laser at 50mW power employing the Andor FRAPPA photomanipulation system. Images were subsequently analyzed using Fiji software and plotted using MATLAB software.

#### Lattice light sheet

The lattice light sheet microscope (LLSM) used in these experiments is housed in the Advanced Imaged Center (AIC) at the Howard Hughes Medical Institute Janelia research campus. The system is configured and operated as previously described (Chen et al., 2014). Briefly, T cells were activated on 5mm round glass coverslips (Warner Instruments, Catalog # CS-5R), plasma-treated and coated with anti-CD3 and ICAM1-extracellular domain, and imaged in culture media at 37°C. Samples were illuminated by lattice light-sheet using 488 nm diode lasers (MPB Communications) through an excitation objective (Special Optics, 0.65 NA, 3.74-mm WD). Fluorescent emission was collected by detection objective (Nikon, CFI Apo LWD 25XW, 1.1 NA), and detected by a sCMOS camera (Hamamatsu Orca Flash 4.0 v2). Acquired data were de-skewed as previously described (Chen et al., 2014) and deconvolved using an iterative Richardson-Lucy algorithm. Point-spread functions for deconvolution were experimentally measured using 200nm tetraspeck beads adhered to 5 mm glass coverslips (T7280, Invitrogen) for 488 excitation wavelength.

#### Traction force measurements

Cell root-mean-square (RMS) tractions at the basal surface were quantified by measuring embedded fluorescent submicrometer particle displacement fields in the gel substrates, following published methods(Poh et al., 2012). Briefly, carboxylated red fluorescent submicron beads (0.2 μm in diameter; F-8810, Life Technologies) were embedded in the hydrogel substrates, the substrates were then covalently functionalized with anti-CD3 (2C11, 10µg/ml) and ICAM-1-12XHis (2µg/ml). Cells were seeded at a concentration of 2 million/ml and allowed to adhere at 37°C for 5 min before the traction force was measured. The un-adhered cells were removed by gentle aspiration with warm culture media before imaging the cells and beads. Fluorescence images of the microbeads were taken using Olympus FV1200 laser scanning confocal system equipped with 30X silicone oil objective (1.05NA), and 550nm laser line, both when the cells were adhered, and again after cells were removed from the substrate using EDTA. Using a custom Matlab code, the displacement of the fluorescent beads at the apical surface of the hydrogel were computed. The RMS traction generated by the cell was calculated based on the displacement field and the known stiffness of the hydrogel substrate. The substrate stiffness of the hydrogel was 100 kPa, prepared using published protocols (Engler et al., 2004; Yeung et al., 2005).

### Brownian Dynamics Model of Active Actin Networks during T-cell Activation

We simulated cytoskeletal networks that consist of actin filaments that can polymerize, depolymerize, and nucleate; actin crosslinking proteins (ACPs) that can bind and unbind in a force-sensitive manner; and active myosin II motors that walk along actin filaments generating tension. The model is based on Brownian dynamics and described in detail in our previous work with no inertia(Kim, 2015; Mak et al., 2016):

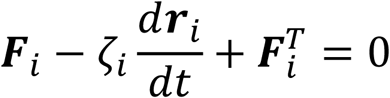

Where, ***Fi*** is the deterministic force determined by the elasticity of the network; *ζi* is the drag of the medium; ***Fi***^***T***-stochastic thermal force; and ***ri*** – is the position of *i*^th^ particle.

The equation is evolved via Euler’s method and solved over time. Extensive details and parameter values, most of which are based on experimental data, of all features in our model can be found here(Mak et al., 2016). Supp. table 1 provides parameter values used in this study in order to investigate the spatial and temporal modulations of the actin cytoskeleton to mimic actin foci formation in the synapse during T cell activation. Cytoskeletal network stresses (in the stress profile plots) are calculated by summing the normal component of tensional forces acting on cytoskeletal components across x-z and y-z cross-sections, divided by the cross-section areas.

To model the T-cell synapse formed against a flat surface, we simulated a thin 3D domain (8 µm × 8 µm × 0.5 µm) with active cytoskeletal components. In this domain, we defined regions with different local kinetics of dynamic cytoskeletal components. At the edges of the cell (400nm width from each edge in the x and y directions), to mimic the integrin-rich peripheral lamella which is important in cell migration, adhesions between actin and the bottom surface are enabled.In the middle of the cell to mimic foci mediated adhesions, a 5 × 5 grid of sites each 400 × 400 nm^2^ enables adhesions between actin filaments and the bottom surface. No boundary adhesions are enabled in other parts of the domain. The nucleation rate of actin filaments is highest at the foci region. To mimic azidoblebbistatin treatment in our simulations, at 100s after the activation of myosin motor walking, we deactivated motor walking and actin-to-substrate adhesions at the top 40% of the domain.

Figure 4A shows the spatial profile of the kinetics of actin nucleation and adhesion used for simulations, and Supp. table 1 shows the parameter values used for the simulations. Because exact rates controlling actin dynamics in live cells are elusive due to the plethora of actin regulating factors (formins, Arp2/3, cofilin, capping proteins, etc.) and spatiotemporal biochemical signaling (e.g. via Rho GTPases), we chose rates of actin turnover comparable to experimental observations (Pollard, 2007) and probed idealized spatial profiles that can capture the spatial distributions of actin seen in T-cells during activation.

### Image analysis and statistics

The images were analyzed using Fiji software. For determination of cell shape (AR), automated cell boundaries were creating by first thresholding the actin images to remove background, watershed plugin to fill the low intensity areas within the cell areas created by threshold operation, and finally with ‘analyze particles’ utility of Fiji. Using the automated boundaries, the raw images were analyzed for the intensity of relevant proteins within each synapse. Foci were estimated as described previously(Kumari et al., 2015). Briefly, a moving window of 1.6×1.6 µm^2^ was used to create a Gaussian blur image of the original raw images. The Gaussian blur image was subtracted from the raw image to create the foci image. Data was plotted using Graphpad Prism or MATLAB. For statistical analysis, Mann-Whitney non-parametric two-tailed test was performed to compare population of cells. In all graphs, unless otherwise mentioned, the dataset was normalized to the mean values of the ‘Control’ population, and then plotted as a ‘normalized’ value. Most graphs highlight the Mean ± SEM in the scatter plot of normalized values, where each dot represents values obtained from a single cell, unless otherwise mentioned.

