## Supplemental for "Cytoskeletal tension actively sustains the migratory T cell synaptic contact"

**Figure S1.** Antigen encounter triggers nucleation of actin foci and cellular symmetry. Related to Figure 1. APS with indicated ligands were incubated with T cells for 2min, fixed, stained with phalloidin-Alexa488 and imaged using SIM. The graph shows quantification of cell shape in cells. Scale bar, 5µm.

**Figure S2.** The aspect ratio (AR) reports on synaptic interface elongation associated with radial interface symmetry breaking. Related to Figure 1. (A) Interference reflection microscopy (IRM) of mouse $CD4^+$ T cells (T cells) shows that significant changes in shape and motility of T cell contact interfaces can be recorded within a time span of 2min, between their 'arrested' (sedentary) and 'motile' states. The motile or arrested cells were manually identified in the time-lapse images, 20 min post their initial contact with APS, and associated mean aspect ratios and speed were analyzed over a time span of 2min. The graph shows an average value of sped or AR spanning 2min. (B) Alteration in speed and shape measured at the population level during synapse breaking. Snapshots of T cells from time-lapse IRM imaging after seeding on APS, with overlaid center-of-mass tracks over time (in color). Shown at right are the speed and aspect ratios calculated within a 2min window of observation at 5 min or 20 min post cell seeding; points are individual cells. Scale bar, 5µm.

**Figure S3**. (A-B) Image processing scheme to utilized to extract and quantify foci on per cell basis. Related to Figure 1. The images show 2D F-actin intensity as marked by phalloidin staining (top images), or a 3D view of spatial distribution of intensities in the phalloidin images (bottom images). To process the raw images for extracting foci intensities from overall F-actin signal, a Gaussian mask was generated by using a 1.6µm X 1.6µm rolling window, as optimized previously[1]. Subtraction of the mask image from the raw image generated a processed image that could be quantified to measure the average intensity contributed by the foci. Note that while this method reliably identifies the foci in raw images and reduces intensity contribution from the non-foci uniform lamellar area, the peripheral lamellipodial network still contributes a background of ~35% to the total foci intensity, regardless of the presence of profuse foci in arrested synapse, or their visible reduction in the motile phase.

**Figure S4.** Calcium sequestration using BAPTA does not predispose cells to synapse breaking. Related to Figure 1. T cells were incubated with APS for 5 min or for 20 min, along with vehicle control or BAPTA and EGTA in the last 10 min of incubation, fixed and imaged using SIM (A). Note that the BAPTA-treated cells retain symmetry more than the control cells, and display significantly more foci, even when they have comparable talin recruitment at the synapse (B).

**Figure S5.** Endogenous pCasL serves as a reliable mechanotransduction marker in T cells. Related to Figure 3. (A) A schematic of mechanosensitive CasL phosphorylation in T cells. CasL is recruited to the signaling TCR clusters via LCK via its Src kinase Binding domain (SB), and interacts with the F-actin cytoskeleton and adhesion complex binding proteins such as FAK via its SH3 domain. Mechanical tension created due to polymerization and remodeling of F-actin at the foci leads to conformational changes in CasL, exposing tyrosine motifs in its substrate domain. These tyrosine residues are phosphorylated by the local Src-Family kinases, and could be immunolabelled to assess TCR-proximal actin cytoskeletal strains. (B, C) Colocalization index of foci and pCasL shows a high degree of association (left plot in C), and correlation with foci intensity per cell (right plot in C). (D) pCasL levels are sensitive to broad actin cytoskeletal perturbation. T cells from WT mice were treated with Latrunculin A (LatA), or left untreated, during incubation with substrate. Cells were subsequently fixed and processed for immunostaining and imaged using TIRF microscopy. ** =0.001 for pCasL. (E) WT T cells show reduced pCasL in polarized synapses. $p$ <0.0001.

**Figure S6.** Related to Figure 3. (A) Initial adhesion and spreading kinetics is comparable in WASP$^{-/-}$ and WT T cells activated on the antigenic surface, WASP$^{-/-}$ cells however show interface elongation at earlier time point (~3 min) than the WT cells. T cells were incubated with substrates on a temperature-controlled microscope stage, imaged live using TIRFM (Movie 5), and analyzed for spreading and shape elongation. (B) Synapse breaking in WT and WASP-/- cells. Images are snapshots of T cells from time-lapse IRM imaging after seeding on APS, with overlaid center-of-mass tracks over time (in color); the graph showed measured speed.

**Figure S7.** Related to Figure 3. TCR engagement-induced phosphorylation of early signaling molecules Zap70 (A), SLP76, LAT (B), and the recruitment of CasL to the synapse in this setting (C).

**Figure S8.** Related to Figure 3. (A) SIM imaging of 2min WASP$^{-/-}$ T cells synapses shows that these cells are able to initially generate radially symmetric ICAM-1 ring in their synapse. Cells were incubated with lipid bilayers reconstituted with anti-CD3 and ICAM1-Cy5, fixed, stained for F-actin and talin and visualized using SIM. (B) Integrin hyperactivation does not rescue symmetry defects in WASP$^{-/-}$ T cells. WT or WASP$^{-/-}$ T cells were incubated with APS in the presence or absence of 0.5mM $MnCl_2$ for 5min, fixed and processed for talin, pCasL and F-actin visualization, and imaged using TIRFM.

**Figure S9.** Related to Figure 3. Intracellular calcium flux is not enough to revert asymmetry in $WASP^{-/-}$ cells. T cells from WT or $WASP^{-/-}$ mice were incubated with APS in the presence of DMSO or 1µM Thapsigargin (Thapsi) for 5'. The cells were then fixed and processed for talin, pCasL and F-actin visualization. Note that while Thapsigargin treatment is unable to restore symmetry in $WASP^{-/-}$ cells (A,B), it induces downregulation of talin in both WT and $WASP^{-/-}$ T cells (quantification in B).

**Figure S10.** Related to Figure 3. (A) pCasL-enriched actin foci in antigen-specific cell-cell conjugate setting. BMDCs loaded with OTII peptide were incubated with mouse WT or $WASP^{-/-}$ OTII transgenic $CD4^{+}$T cells for 5', fixed and processed for SIM imaging. The image shows maximum intensity projection from 2µm depth of the synaptic area of a single T cell, marked by a white box. The graph on the right shows the intensity profiles of F-actin and pCasL across a single foci, outlined in the adjacent image in the 'WT' case. (B) Representative Total Internal Reflection Fluorescence microscopy (TIRF) images of primary human $CD4^{+}$ T cells isolated from healthy individuals or WAS patients and activated for 5' on APS. Graph on the right shows quantification of actin foci, pCASL, and cell AR normalized to values of control healthy individual cells.

**Figure S11.** Related to Figure 3. Transient reduction of WASP levels in human cells elicits defects comparable to those observed in murine T cells genetically deficient of WASP. (A) Human $CD4^{+}$ T cells transduced with control lentivirus or with lentivirus delivering WASP shRNA (as described in Kumari et. al., eLife, 2015), seeded onto superantigen-loaded HUVEC cells for 5', then fixed and imaged using SIM superresolution imaging. (B) Foci polymerization role of WASP underlies its mechanical tension-generating activity. Human $CD4^{+}$T cells were transfected with human WT WASP-GFP, WASPΔC or with WASP shRNA (shR)-transducing lentiviral particles. The data shoes that the WASP shR and WASPΔC reduce foci and pCasL at the synapse to a similar extent. The remaining foci in the cells in the shR and WASPΔC are contributed by APC cytoskeletal features underneath the synapse, which are quantified along with foci by our foci extraction algorithm outlined in Figure S3.

**Figure S12.** Related to Figure 5. Endogenous myosinII distribution at the synapse in the stable (upper panels) and broken phase (lower panels). Arrows shows the location of myosin puncta juxtaposed with the actin foci.

#### Videos:

**Video S1. Related to Figure 1.** Six different examples of cells breaking their sedentary primary contacts and showing interface shape elongation and a shift in motility, imaged using IRM, indicate that rapid shape transitions can be quantified with 2 min of time duration.

**Video S2. Related to Figure 1.** IRM live imaging of T cells using reveals a significant shift away from the primary synapse site within 20 min of APS encounter. The images were negatively contrasted to identify cell positions and better highlight individual cell boundaries using automated cell outlining (object identification) routine in ImageJ. The residual material left by the T cells on the primary contact site is reminiscent of membrane fragments, as described in[5].

**Video S3. Related to Figure 2.** LLSM live imaging of mouse T cell synapse expressing LifeAct-GFP, during transition to the motile phase.

**Video S4. Related to Figure 2.** LifeAct-GFP expressing WT or WASP-/- T cells reveal differential dynamics of lamella vs. the foci. Cells were allowed to attach to the APS for 5min (t=0 in the movie), and then imaged using TIRFM.

**Video S5. Related to Figure 3.** $WASP^{-/-}$ cells break contact symmetry faster than the WT T cells. T cells isolated from WT or $WASP^{-/-}$ mice were imaged live using IRM for 10 min at 3 frames/min. The images were negatively contrasted for automated boundary identification (Movie 8; see 'Methods') and tracked for the center of mass movement. Bottom panels represent the cell traces overlaid in top panels.

**Videos S6-9**. **Related to Figure 4.** Simulations showing evolution of F-actin network in the WT (whole stable synapse, Movie 6; magnified view of cytoskeletal dynamics around foci, Movie 7), $WASP^{-/-}$ (predisposed to breaking, Movie 8), and a WT synapse transitioning into polarized state (Movie 9), soon after the initial antigen encounter and spreading. Each movie represents a synapse as a rectangular simulation space, based on the scheme presented in Figure 4A. Note that WT synapse in Movie 9 loses established symmetry the instant WASP is removed from the simulation.

**Video S10. Related to Figure 5.** LLSM imaging of cells treated with CK666 show a loss of foci and altered actin dynamics in the synapse similar to that in the case of $WASP^{-/-}$ cells.

**Video S11**. **Related to Figure 5**. Simulation showing the effect of localized myosin perturbation on F-actin network connectivity on the tension within the synapse. The simulation scheme described in Figure 4A was used to create F-actin distribution in synapse, where a perturbation in myosinII was introduced in a rectangular shaped subsynaptic region, shortly after the synapse was established (corresponding to the image shown in Figure 5h).

**Video S12. Related to Figure 6.** Control or Azidoblebb.-treated cells, corresponding to the images shown in Figure 4h (photoactivable inhibition of myosinII).

### Figure S1

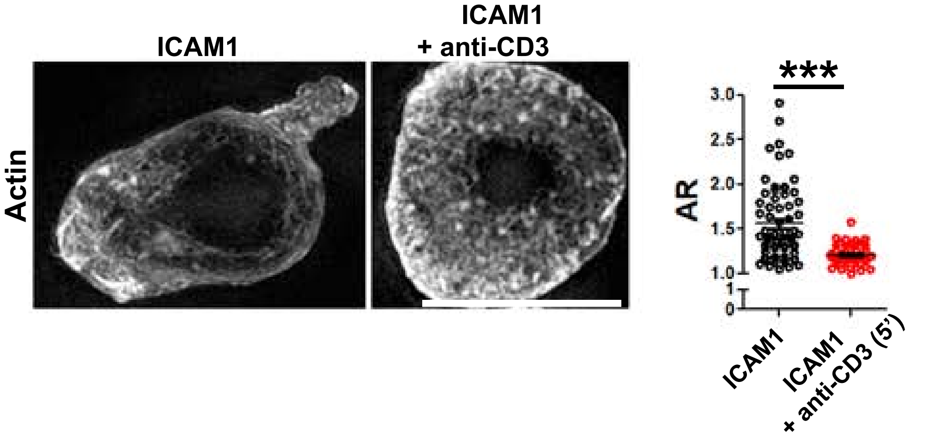

ICAM1
ICAM1
+ anti-CD3
Actin
***
3.0
2.5
2.0
1.5
1.0
1
0
AR
ICAM1
ICAM1
+ anti-CD3 (5')

### Figure S2

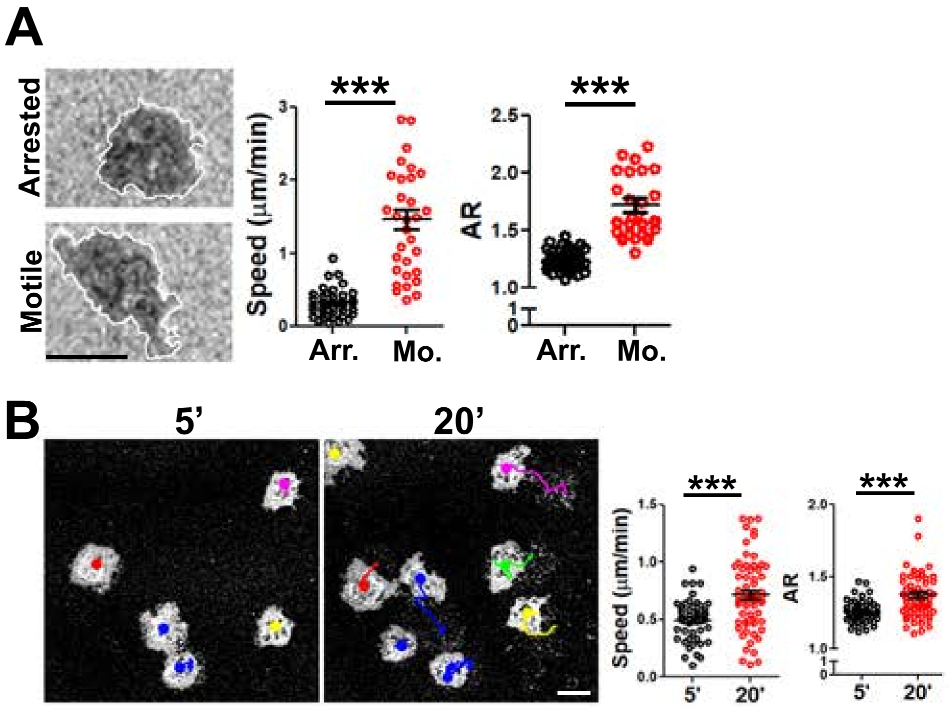
A
Arrested
Motile
***
Speed (μm/min)
Arr.
Mo.
***
AR
2.5
2.0
1.5
1.0
Arr.
Mo.
B
5'
20'
***
Speed (μm/min)
5'
20'
***
AR
5'
20'

### Figure S3

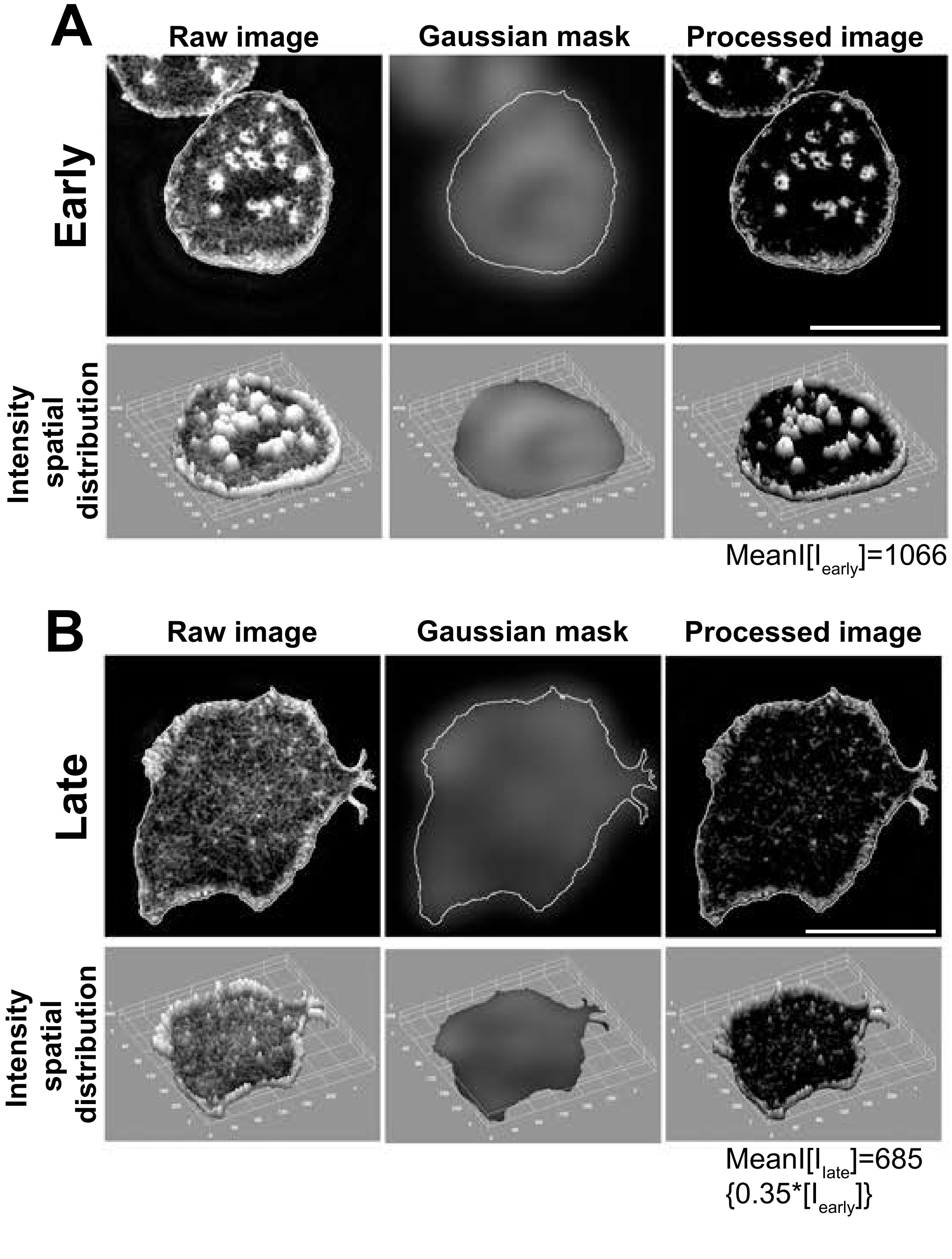
A
Raw image
Gaussian mask
Processed image
Early
Intensity spatial distribution
MeanI[I$_{early}$]=1066
B
Raw image
Gaussian mask
Processed image
Late
Intensity spatial distribution
MeanI[I$_{late}$]=685
{0.35*[I$_{early}$]}

### Figure S4

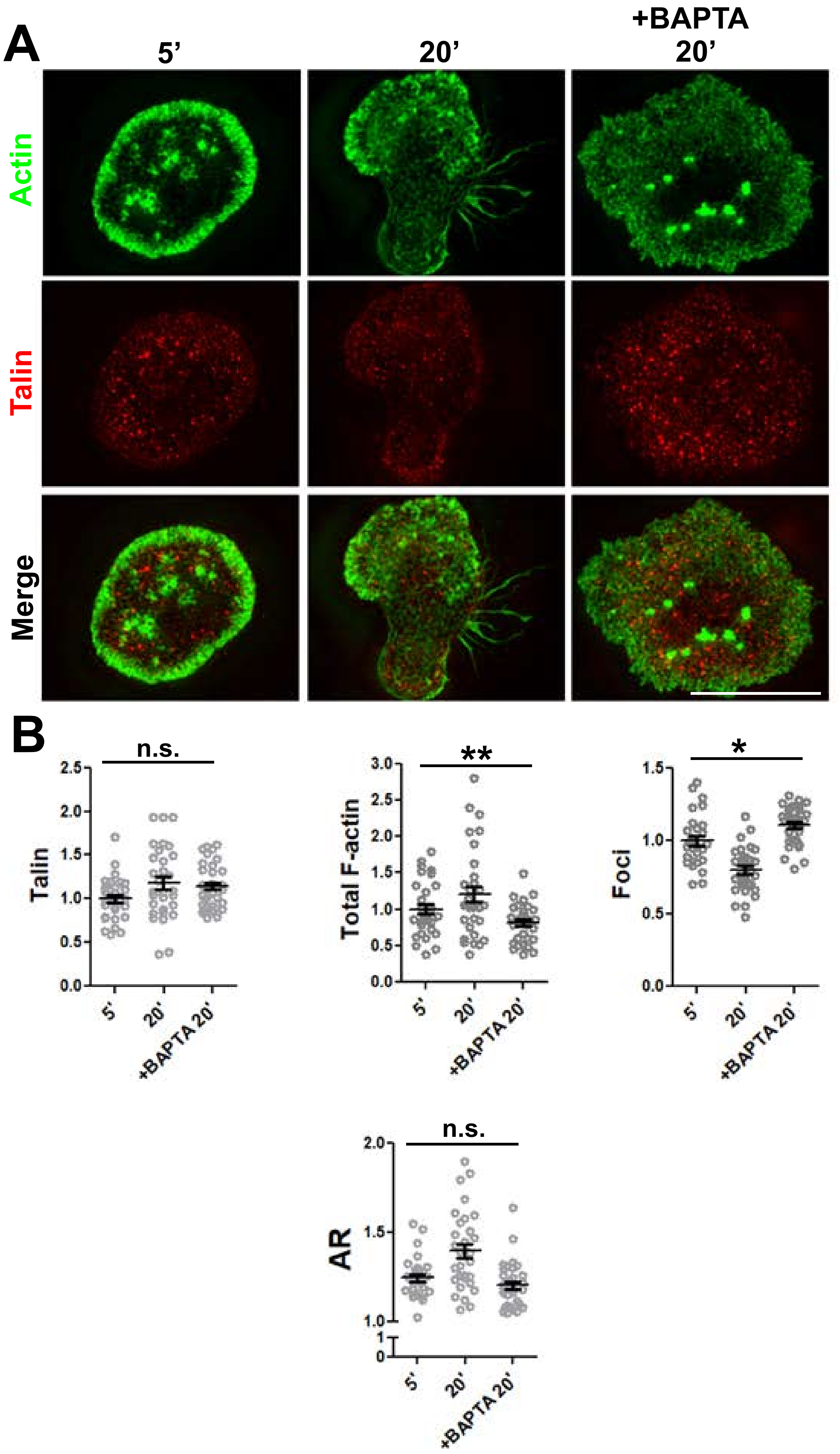
A
5'
20'
+BAPTA 20'
Actin
Talin
Merge
B
n.s.
Talin
**
Total F-actin
*
Foci
n.s.
AR
5'
20'
+BAPTA 20'

### Figure S5

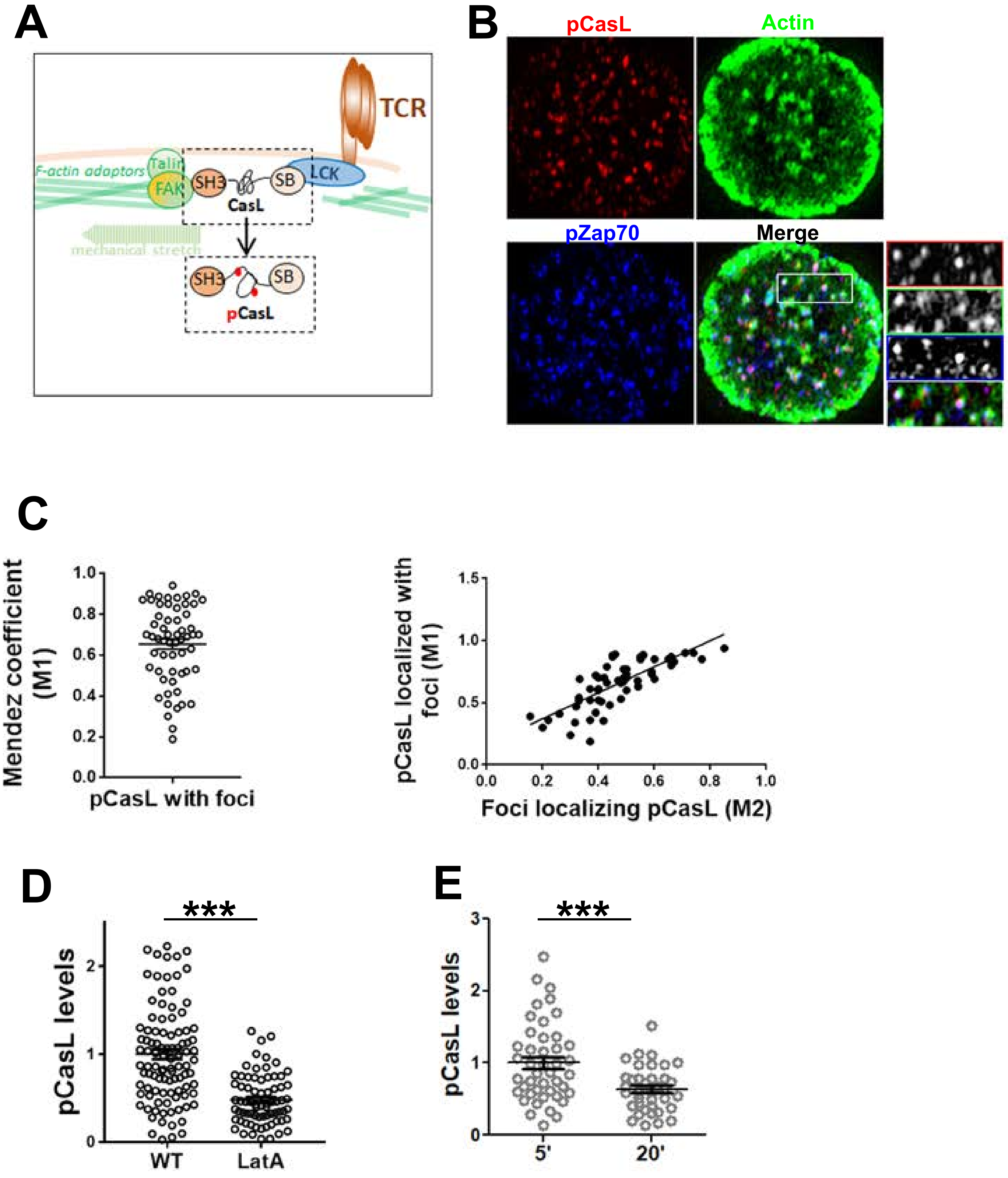
A
TCR
F-actin adaptors
Talin
FAK
SH3
SB
LCK
CasL
mechanical stretch
pCasL
B
pCasL
Actin
pZap70
Merge
C
Mendez coefficient (M1)
pCasL with foci
pCasL localized with foci (M1)
Foci localizing pCasL (M2)
D
***
pCasL levels
WT
LatA
E
***
pCasL levels
5'
20'

### Figure S6

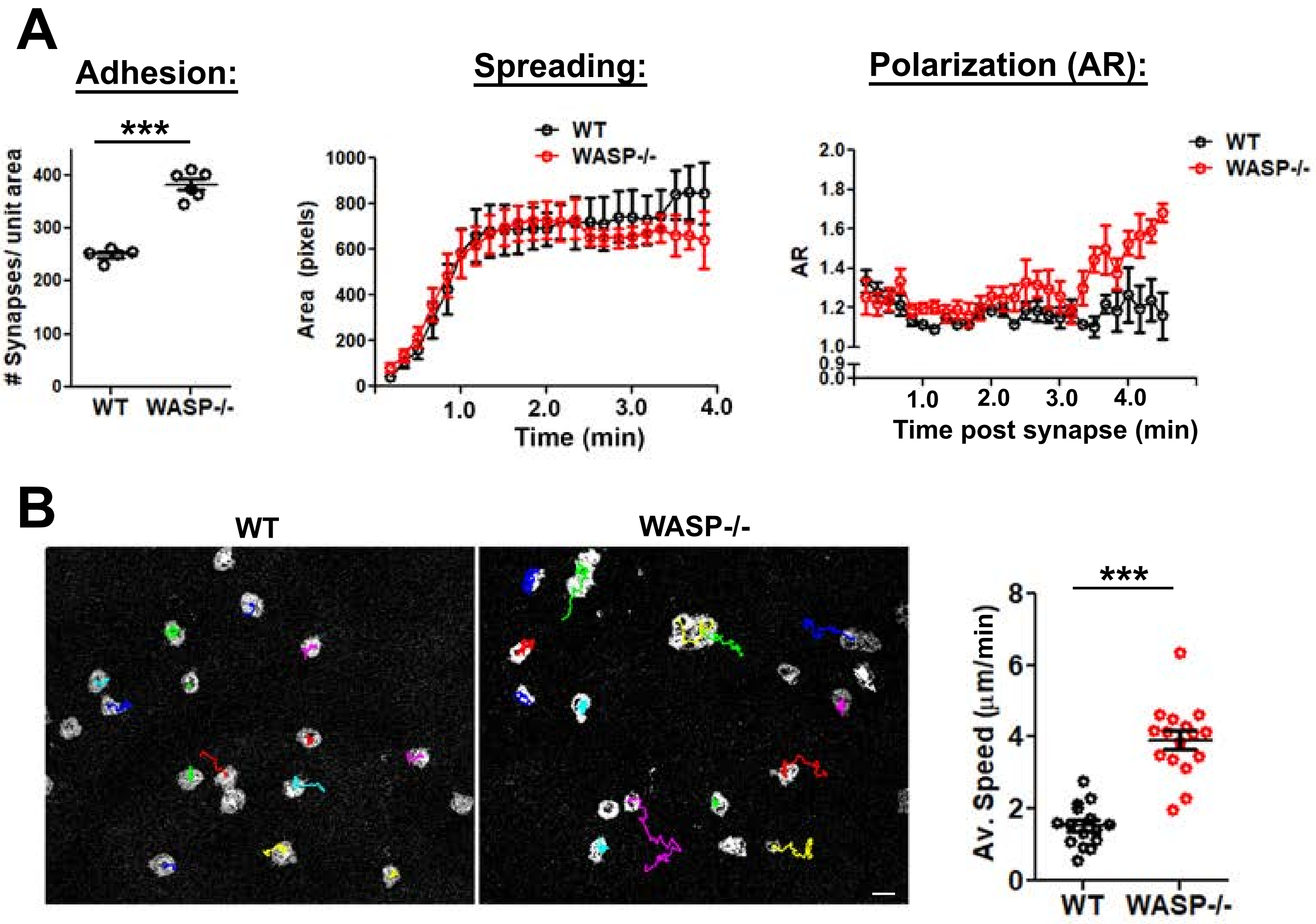
A
Adhesion:
***
### Synapses/ unit area
0
100
200
300
400
WT
WASP-/-
Spreading:
WT
WASP-/-
Area (pixels)
0
200
400
600
800
1000
1.0
2.0
3.0
4.0
Time (min)
Polarization (AR):
WT
WASP-/-
AR
0.0
0.9
1.0
1.2
1.4
1.6
1.8
2.0
1.0
2.0
3.0
4.0
Time post synapse (min)
B
WT
WASP-/-
***
Av. Speed (μm/min)
0
2
4
6
8
WT
WASP-/-

### Figure S7

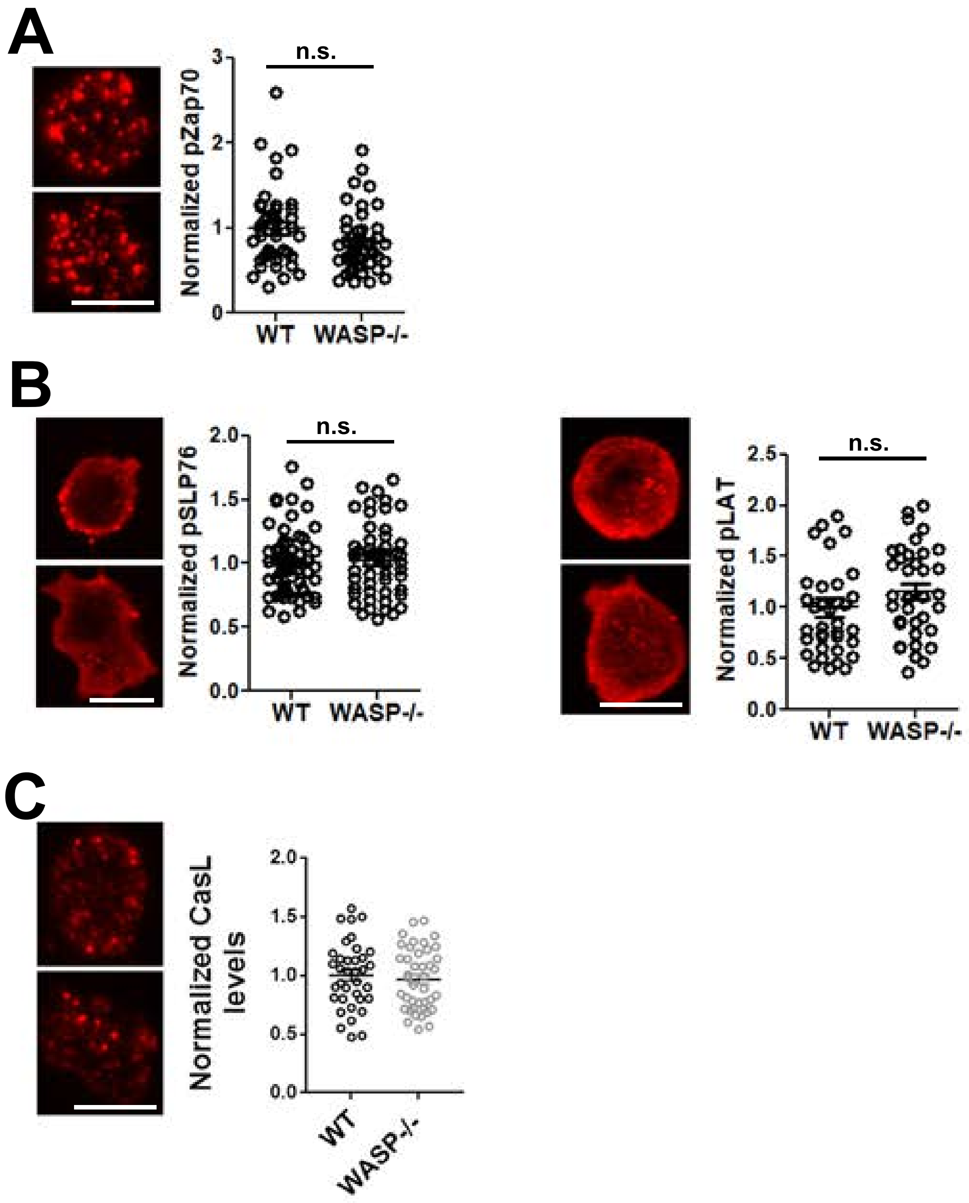

A
n.s.
Normalized pZap70
0
1
2
3
WT
WASP-/-
B
n.s.
Normalized pSLP76
0.0
0.5
1.0
1.5
2.0
WT
WASP-/-
n.s.
Normalized pLAT
0.0
0.5
1.0
1.5
2.0
2.5
WT
WASP-/-
C
Normalized CasL levels
0.0
0.5
1.0
1.5
2.0
WT
WASP-/-

### Figure S8

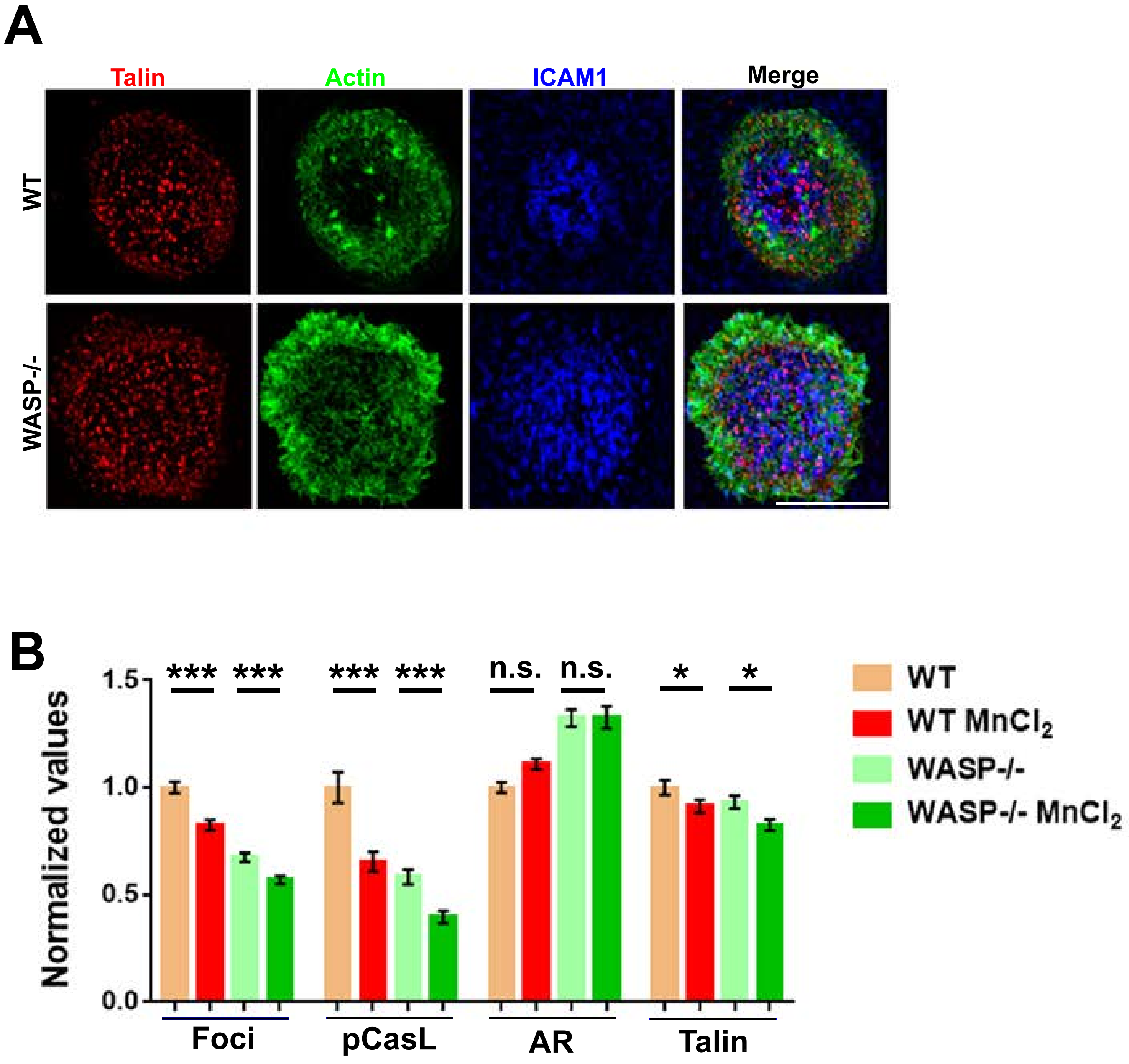
A
Talin
Actin
ICAM1
Merge
WT
WASP-/-
B
Normalized values
1.5
1.0
0.5
0.0
***
***
***
***
n.s.
n.s.
*
*
Foci
pCasL
AR
Talin
WT
WT MnCl2
WASP-/-
WASP-/- MnCl2

### Figure S9

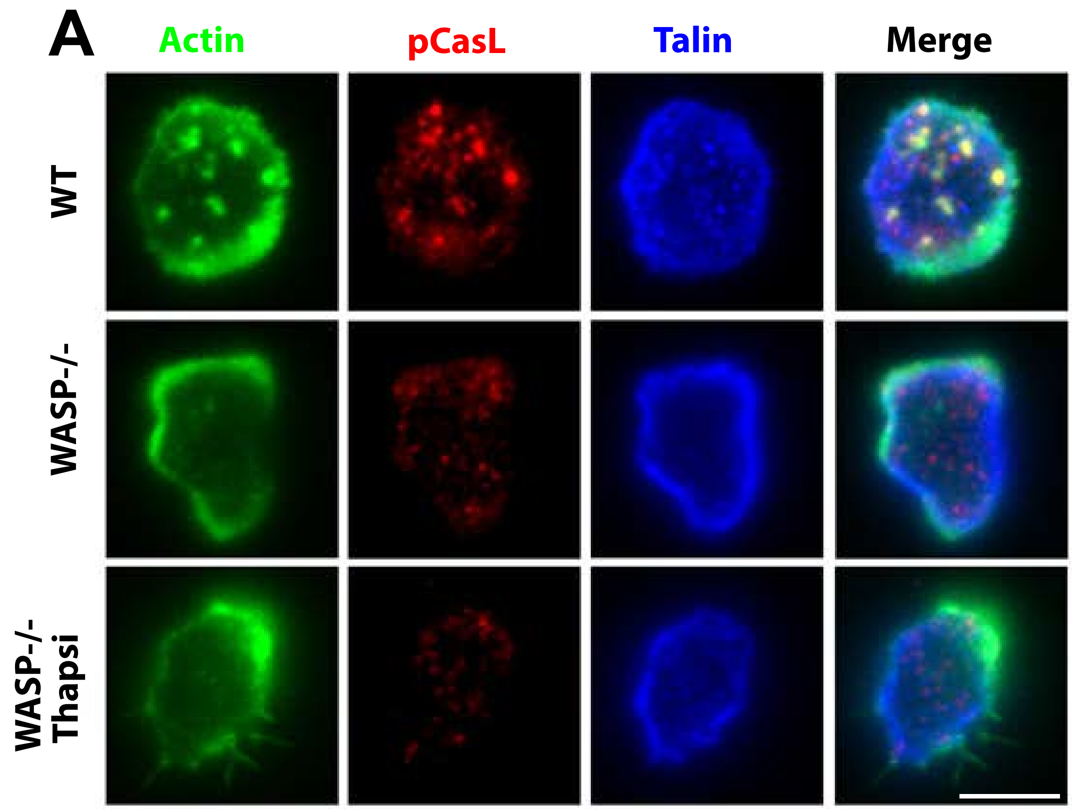

A
Actin
pCasL
Talin
Merge
WT
WASP-/-
WASP-/- Thapsi


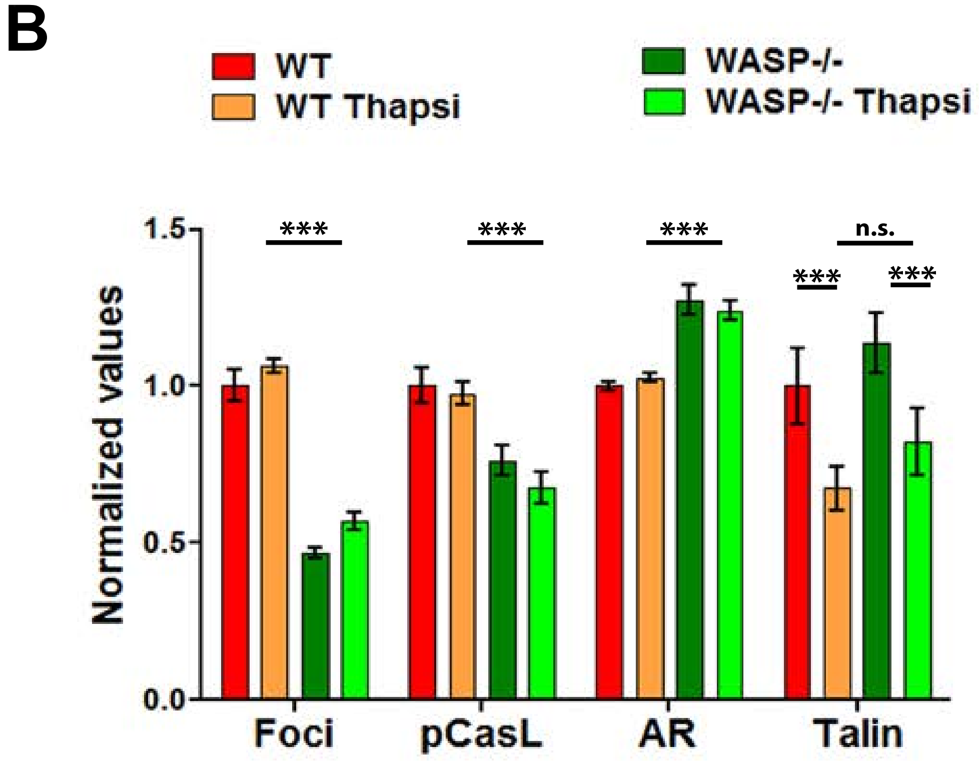

B
WT
WT Thapsi
WASP-/-
WASP-/- Thapsi
Normalized values
1.5
1.0
0.5
0.0
***
***
***
n.s.
***
***
Foci
pCasL
AR
Talin

### Figure S10

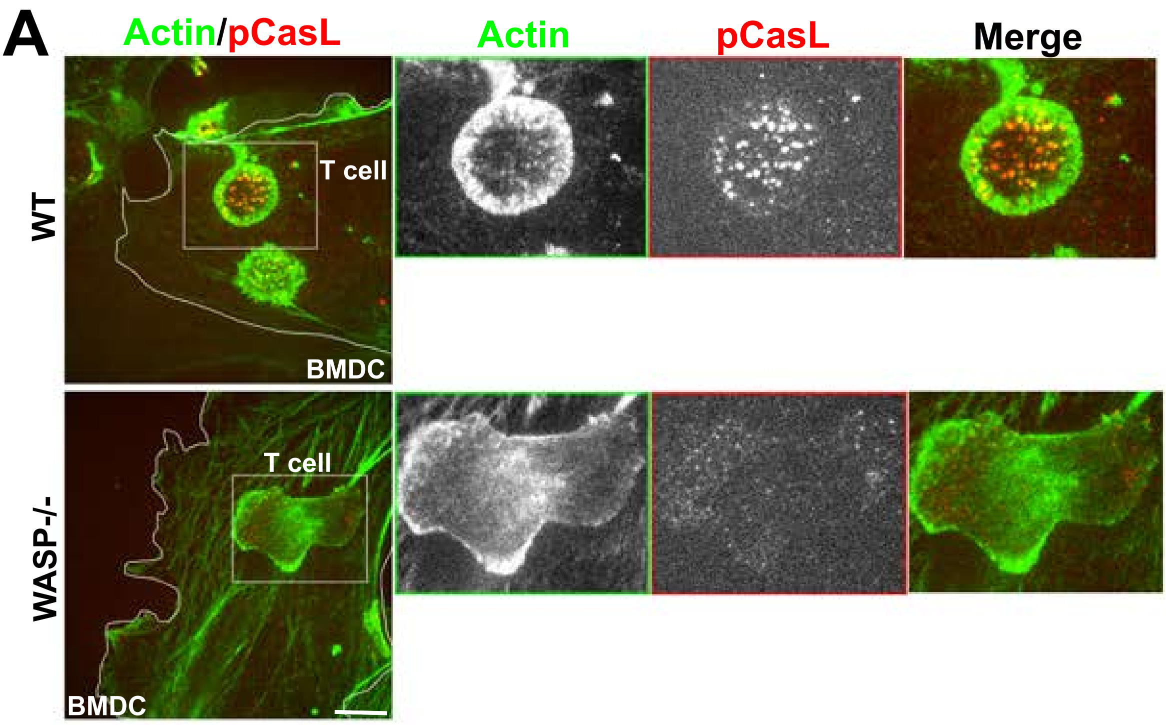

A
Actin/pCasL
Actin
pCasL
Merge
WT
T cell
BMDC
WASP-/-
T cell
BMDC


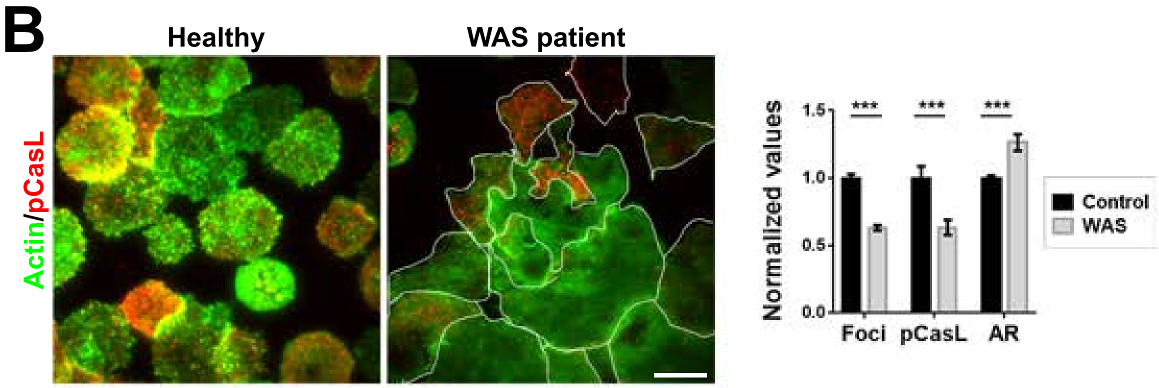

B
Healthy
WAS patient
Actin/pCasL
Normalized values
1.5
1.0
0.5
0.0
***
***
***
Foci
pCasL
AR
Control
WAS

### Figure S11

A

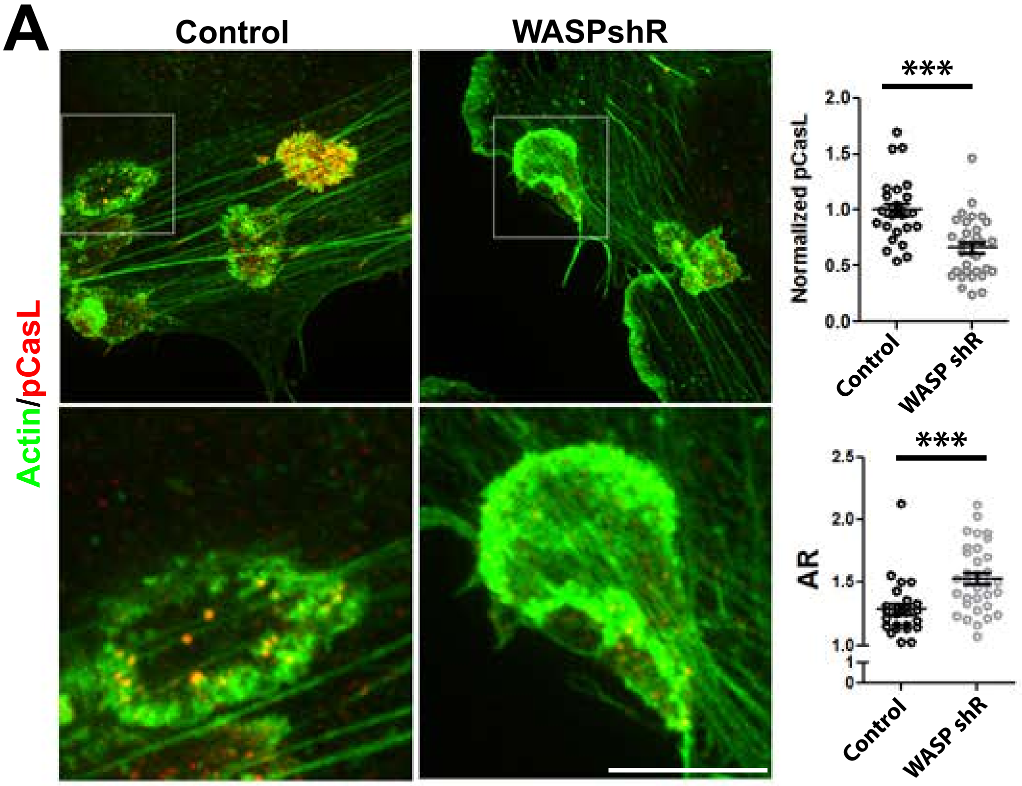


B

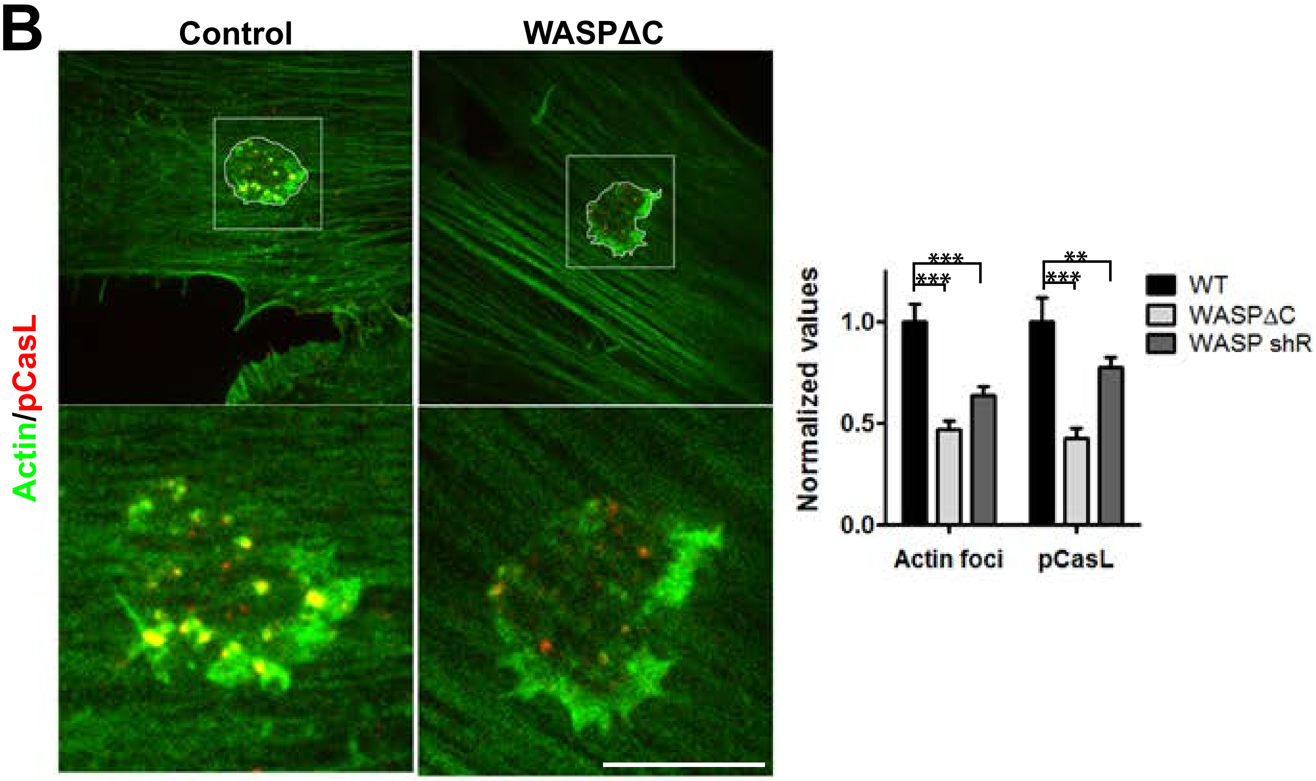

### Figure S12

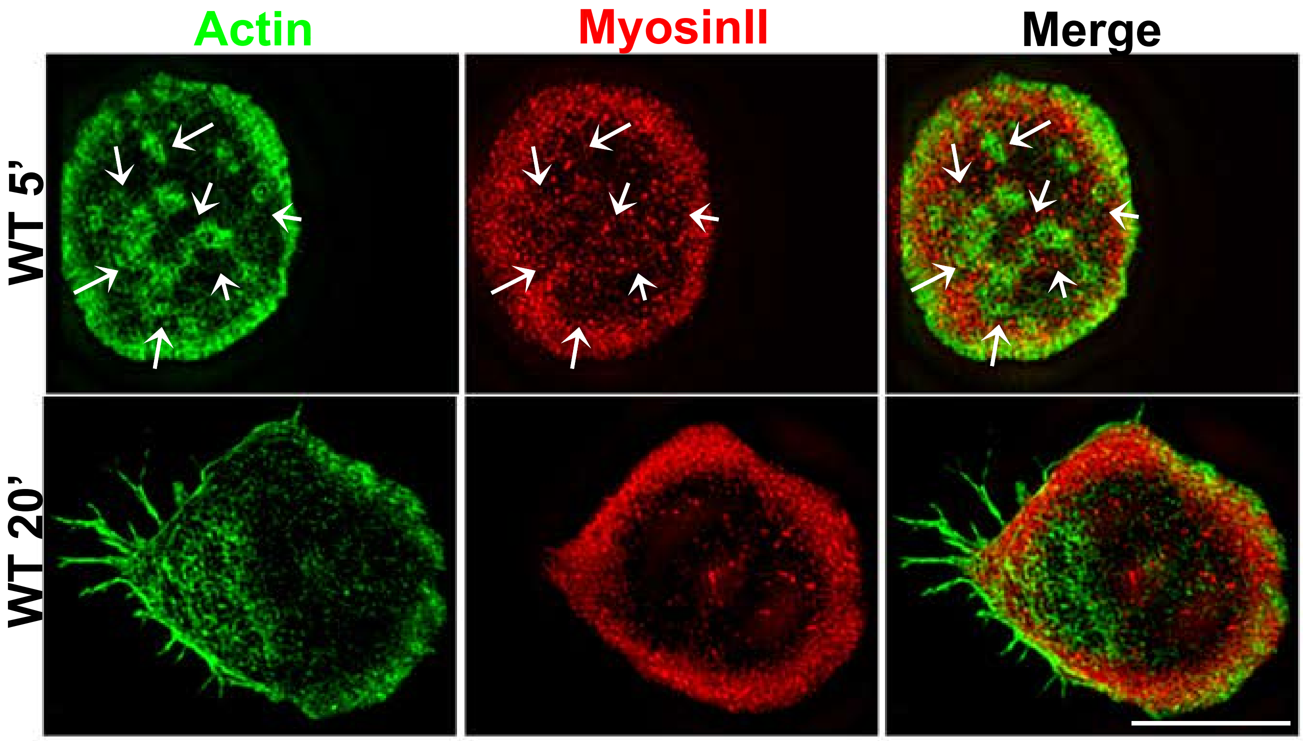

**Table S1. Related to Figure 4:** Model parameters. Parameter values are similar to those used in our previous work where the entire parameter list can be found (*1*, *2*). This table shows values important and/or tuned in this study.

| Symbol | Definition | Value |
|---|---|---|
| $C_{\mathrm{A}}$ | Actin concentration | 25 [μM] |
| % Motors | [Ratio of $C_{\mathrm{M}}$ to $C_{\mathrm{A}}$] × 100 | 1 |
| % ACPs | [Ratio of $C_{\mathrm{ACP}}$ to $C_{\mathrm{A}}$] × 100 | 1 |
| $k^0_{\mathrm{u,ACP}}$ | Zero-force unbinding rate coefficient of ACP | 0.115 [$s^{-1}$](*3*) |
| $\lambda_{\mathrm{u,ACP}}$ | Compliance of a bond for ACP unbinding | $1.04\times10^{-10}$ [m](*3*) |
| $k_{a,p}$ | Actin polymerization rate | 0.3 [$\mu M^{-1}$ $s^{-1}$] |
| $k_{a,d}$ | Actin depolymerization rate | 0.3 [$s^{-1}$] |
| $k_{a,n}$ | Actin nucleation rate | $n * 0.001$ [$\mu M^{-1}$ $s^{-1}$] |
| $n$ | Actin turnover scaling factor | 1, at foci<br>0.01, at edges |
| $k_{adh,unb}$ | Zero-force unbinding rate of adhesions between actin and substrate | $n_{adh} * k^0_{u,ACP}$ |
| $\lambda_{u,adh}$ | Compliance of a bond for adhesions | $\lambda_{u,ACP}$ |
| $n_{adh}$ | Adhesion unbinding scaling factor | 0.1, at foci<br>10, at edges |

Note: The bonds linking actin filaments with actin filaments (ACPs) and actin filaments with the substrate (adhesions) are modeled as slip bonds via Bell's equation (*4*) with unbinding rates equal to:

$$k_u = k_u^0 \exp\left(\frac{\lambda_u |F|}{k_B T}\right)$$

where $|F|$ is the tension acting on the bond, $k_B$ is the Boltzmann constant, and $T$ is temperature.


1. M. P. M. Wonyeong Jung, Taeyoon Kim, F-actin cross-linking enhances the stability of force generation in disordered actomyosin networks. *Computational Particle Mechanics* **2**, 317 (2015).

2. M. Mak, M. H. Zaman, R. D. Kamm, T. Kim, Interplay of active processes modulates tension and drives phase transition in self-renewing, motor-driven cytoskeletal networks. *Nature communications* **7**, 10323 (Jan 8, 2016).
3. J. M. Ferrer *et al.*, Measuring molecular rupture forces between single actin filaments and actin-binding proteins. *Proceedings of the National Academy of Sciences of the United States of America* **105**, 9221 (Jul 8, 2008).
4. G. I. Bell, Models for the specific adhesion of cells to cells. *Science* **200**, 618 (May 12, 1978).
